# Predicting Biological Age Using an Accumulated Neurotoxicity Biomarker for Amyloid Beta Oligomers

**DOI:** 10.1101/2025.02.28.640920

**Authors:** Andrey V. Kuznetsov

## Abstract

This study proposes using accumulated neurotoxicity, defined as the time integral of Aβ oligomer concentration, as a biomarker for neuronal aging. A relationship between biological age and accumulated neurotoxicity is proposed. Numerical analysis guided the development of a new analytical solution linking the biological and calendar ages of neurons. The effects of Aβ monomer and oligomer half-lives—key indicators of proteolytic efficiency—on biological age are examined. Both constant and age-dependent (exponentially increasing) half-life scenarios are considered. The findings indicate that increasing the half-life of Aβ monomers and oligomers with age accelerates biological aging. Reducing Aβ monomer production is shown to slow biological aging, with a linear relationship established between these two quantities. Additionally, biological age is found to depend linearly on the half-deposition time of Aβ oligomers into senile plaques. The model demonstrates that biological age is irreversible, providing a theoretical explanation for why plaque-clearing therapies cannot reverse established cognitive impairment. The model also demonstrates that biological age is path-dependent rather than state-dependent.

## 1. Introduction

Aging is a universal process in complex multicellular organisms. Understanding aging is, therefore, critically important. It seems logical to associate aging with irreversible changes that occur in an organism as its chronological age increases. However, the term “irreversible” should be used cautiously. In thermodynamics it is used for a process in which total entropy generation, including the system and its surroundings, is positive. Although nearly all biological processes are thermodynamically irreversible, this does not imply a continuous increase in an organism’s entropy. As open systems, living organisms maintain their low-entropy state by exchanging energy and increasing the entropy of their surroundings.

Reed *et al*. (2024) proposed the term *resilience* to describe a system’s capacity to return to its original state following a perturbation. In the context of aging, resilience can be understood as the ability to recover from age-related damage. The hydra, a simple organism, appears to possess virtually unlimited resilience, as it is capable of indefinite stem cell renewal and exhibits negligible senescence (Klimovich *et al*., 2018). In contrast, more complex organisms have a finite capacity for self-repair. Many types of damage cannot be fully repaired and accumulate over time, including epigenetic changes (Wang *et al*., 2022), errors in transcription and translation (Vermulst *et al*., 2015), damage caused by reactive oxygen species (Labunskyy & Gladyshev, 2013; Giorgi *et al*., 2018), and various other forms of deterioration. Aging can therefore be understood as a consequence of the biological system’s limited resilience.

Aging arises from the accumulation of damage across multiple biological levels, from molecules to entire organ systems, which the body cannot fully repair. Here, “damage” refers to detrimental changes associated with aging. However, some changes may aid in adaptation to aging, while others may be neutral and have no harmful effects on the organism (Gladyshev *et al*., 2021). When aging is viewed within the organism’s functional domain, certain brain functions may, in some cases of healthy aging, even improve with chronological age as a compensatory mechanism for the decline in overall brain performance (Knights *et al*., 2024).

Studying interventions aimed at slowing aging requires measuring how rapidly an individual is aging. An important factor that influences this rate is biological age, which—unlike chronological age—varies across individuals due to differences in the pace of aging. Various aging clocks, which are computational models based on biomarkers of aging, are used to estimate biological age (Min *et al*., 2024). Examples include epigenetic clocks that analyze DNA methylation patterns, telomere clocks that measure telomere length, and epitranscriptomic clocks that track RNA methylation (Wang *et al*., 2023; Han, 2024; Crimmins *et al*., 2024).

The abundance of aging clocks reflects the complexity of the aging process, which involves numerous biological mechanisms. (Dormann & Lemke, 2024) proposed developing a novel aging clock based on the aggregation of intrinsically disordered proteins (IDPs) such as amyloid beta (Aβ), tau, α-synuclein, and TDP-43. IDPs have a high tendency to aggregate due to the lower free energy of their aggregated state compared to their native, disordered form (Vetri & Foderà, 2015). The significance of an IDP-based aging clock lies in the fact that aging is the primary risk factor for neurodegenerative diseases, including Alzheimer’s (AD) and Parkinson’s, which are closely linked to IDP aggregation.

There is considerable scientific interest in the mechanisms underlying brain aging (Antal *et al*., 2025). This current paper proposes using the accumulated neurotoxicity of Aβ oligomers as a biomarker for the biological age of neural cells (hereafter referred to as biological age). Numerical simulations are conducted to explore the relationship between the proposed biomarker, calendar age, and other model parameters. A hypothetical 70-year timespan is considered.

Unlike the results in Kuznetsov (2025a), which provided analytical solutions only for the limiting cases of infinitely slow (*θ*_1/ 2, *B*_ → ∞) and infinitely fast (*θ*_1/ 2, *B*_ → 0) deposition rates of Aβ aggregates into senile plaques—where *θ*_1/2,*B*_ denotes the time required for half of the free Aβ aggregates to be incorporated into a plaque—this study presents an analytical solution for finite, physiologically relevant half-deposition times of Aβ oligomers into senile plaques. Due to the non-linearity of the governing equations, the behavior of the analytical solution presented in this paper differs significantly from previously reported solutions. The sensitivity of the predicted biological age to model parameters characterizing the efficiency of protein degradation machinery and the Aβ monomer production rate is investigated.

## 2. Materials and models

### 2.1. Mathematical framework for Aβ peptide aggregation

As in Kuznetsov (2025a), Kuznetsov (2025b), the aggregation of Aβ peptides is described using the Finke-Watzky (F-W) model, which simplifies the process into two pseudo-elementary steps: nucleation and autocatalytic growth. During nucleation, initial aggregates form at a continuous rate, while in the autocatalysis phase, pre-existing aggregates catalyze the conversion of monomers into additional aggregates (Morris *et al*., 2008; Iashchishyn *et al*., 2017). These two reaction mechanisms are mathematically represented as follows:

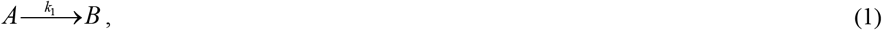

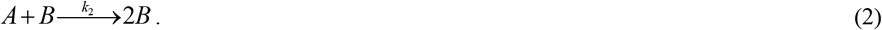

In Eqs. (1) and (2), *A* represents a monomeric peptide, while *B* denotes an amyloid-converted peptide that remains free and has not yet been incorporated into a plaque. The kinetic constants *k*_1_ and *k*_2_ correspond to the nucleation and autocatalytic growth rates, respectively (Morris *et al*., 2008). Primary nucleation, simulated by Eq. (1), involves only monomers, whereas secondary nucleation, described by Eq. (2), requires interactions between monomers and pre-existing free aggregates of the same peptide (Thacker *et al*., 2023).

Conservation equations for Aβ monomers, oligomers, and plaques are formulated for a control volume (CV) illustrated in Fig. 1, situated within a region of the brain where Aβ plaque formation occurs. Under conditions of rapid Aβ monomer diffusion, their concentration ( *C*_*A*_) can be considered nearly uniform throughout the CV. It is further assumed that each CV within this region maintains homogeneous and uniform concentrations of Aβ monomers, oligomers, and plaques. Under this assumption, time *t* serves as the only independent variable in the model. The dependent variables used in the analysis are summarized in Table 1, while the model parameters are summarized in Table 2.

**Table 1.** List of dependent variables used in the model.

| Symbol | Definition | Units |
| --- | --- | --- |
| $C_A$ | Concentration of Aβ monomers | $\mu\text{M}$ |
| $C_B$ | Concentration of free Aβ aggregates (such as oligomers and protofibrils) | $\mu\text{M}$ |
| $C_D$ | Concentration of Aβ aggregates incorporated into amyloid plaques | $\mu\text{M}$ |

**Table 2.**
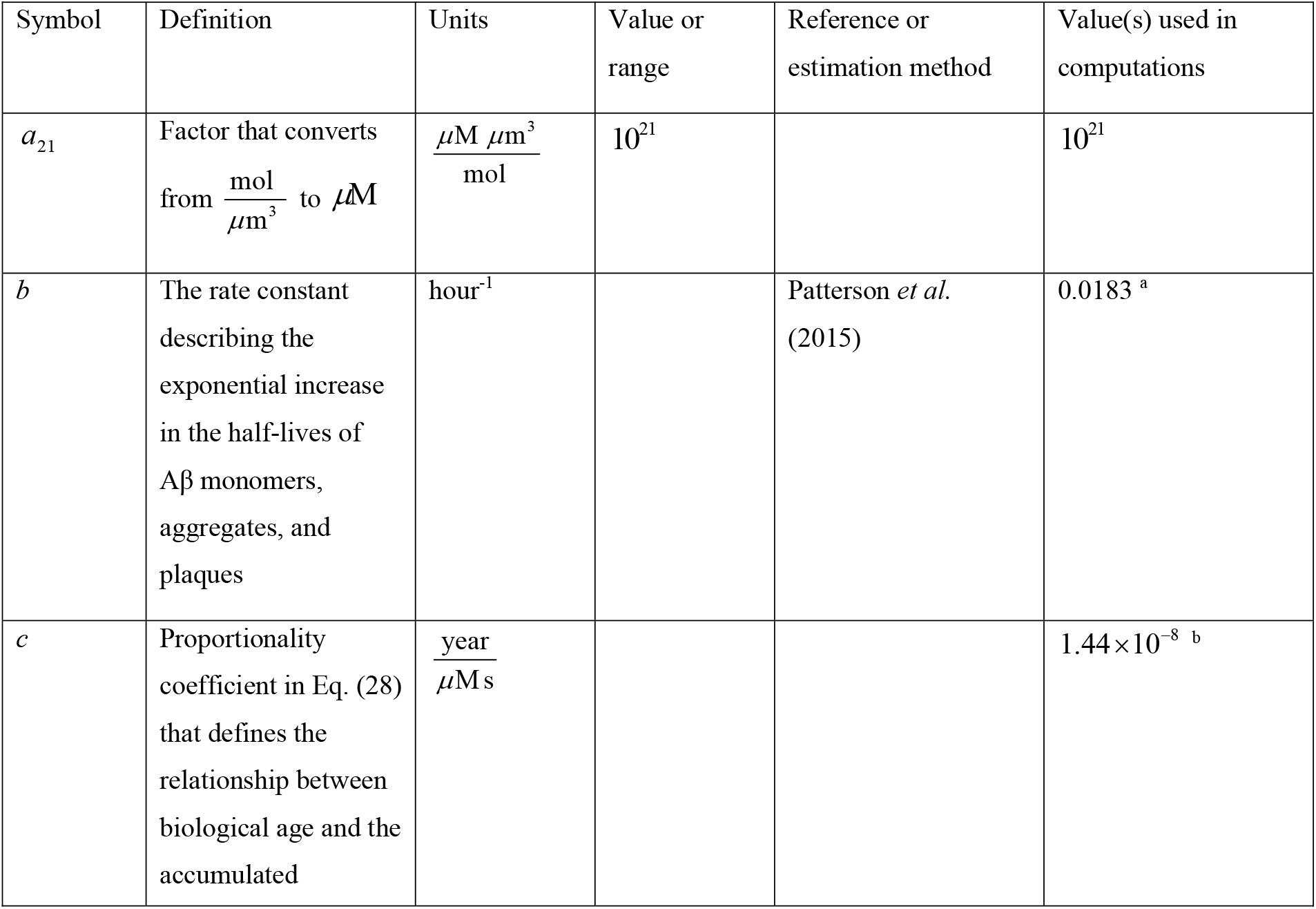
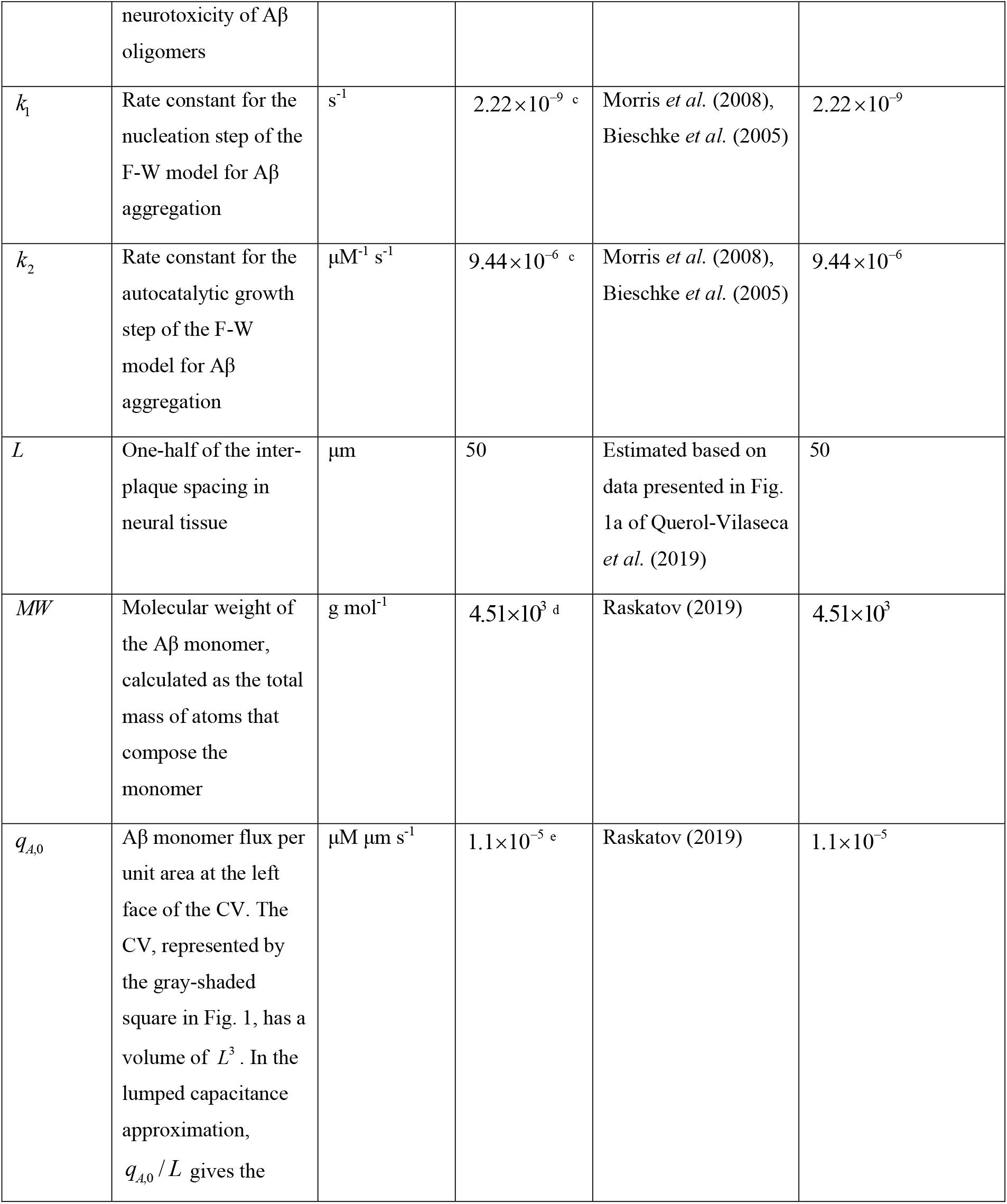
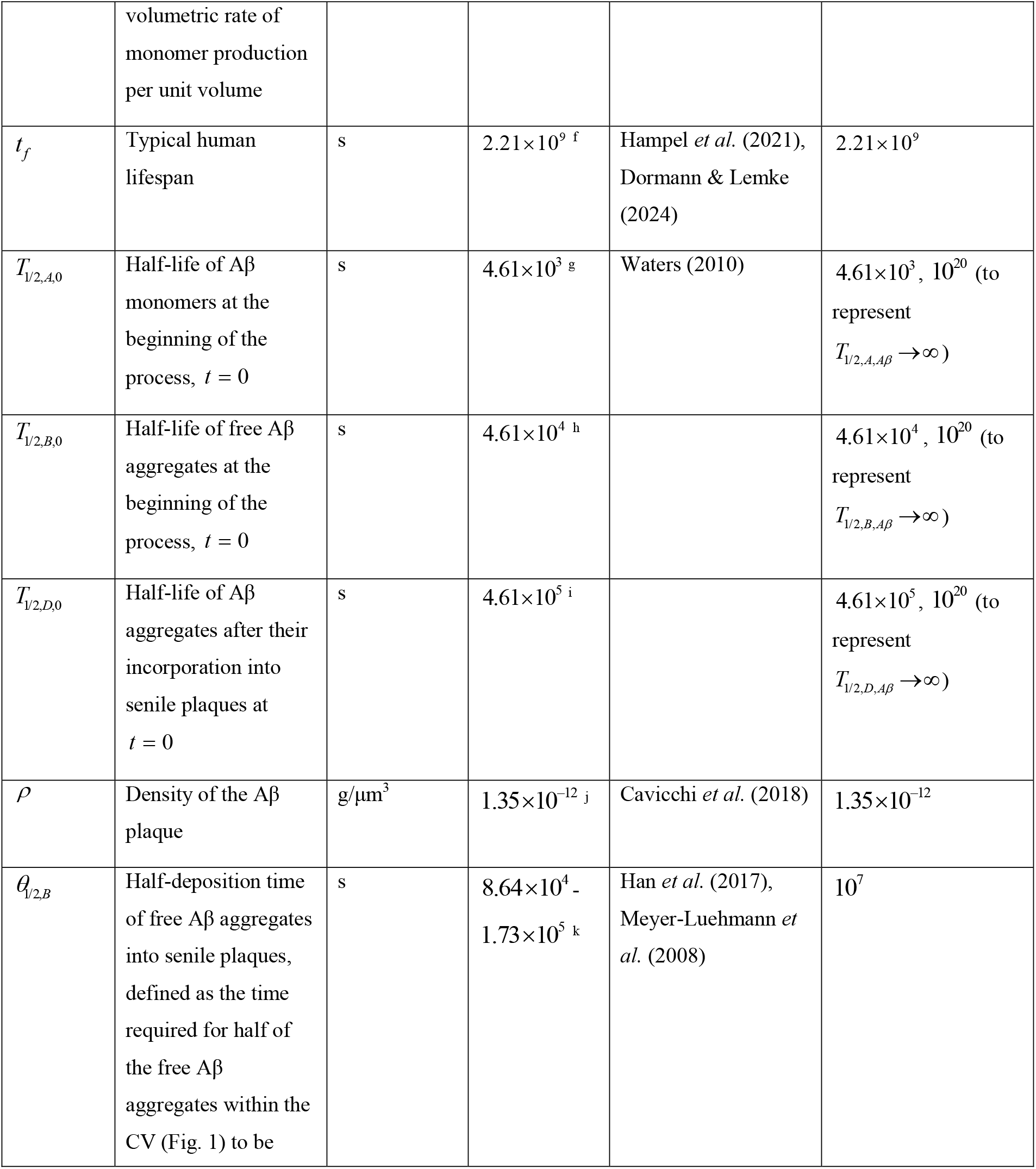
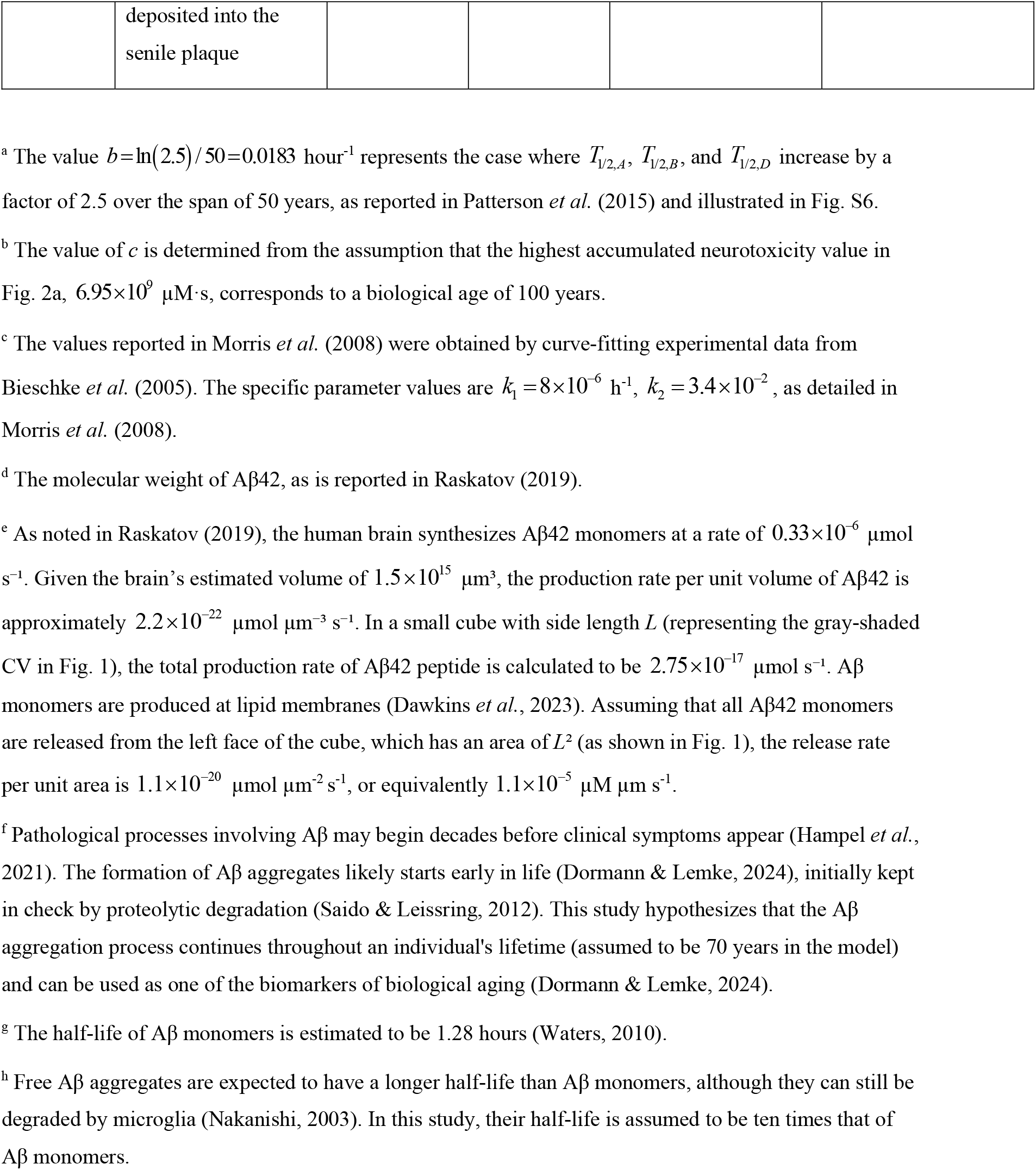
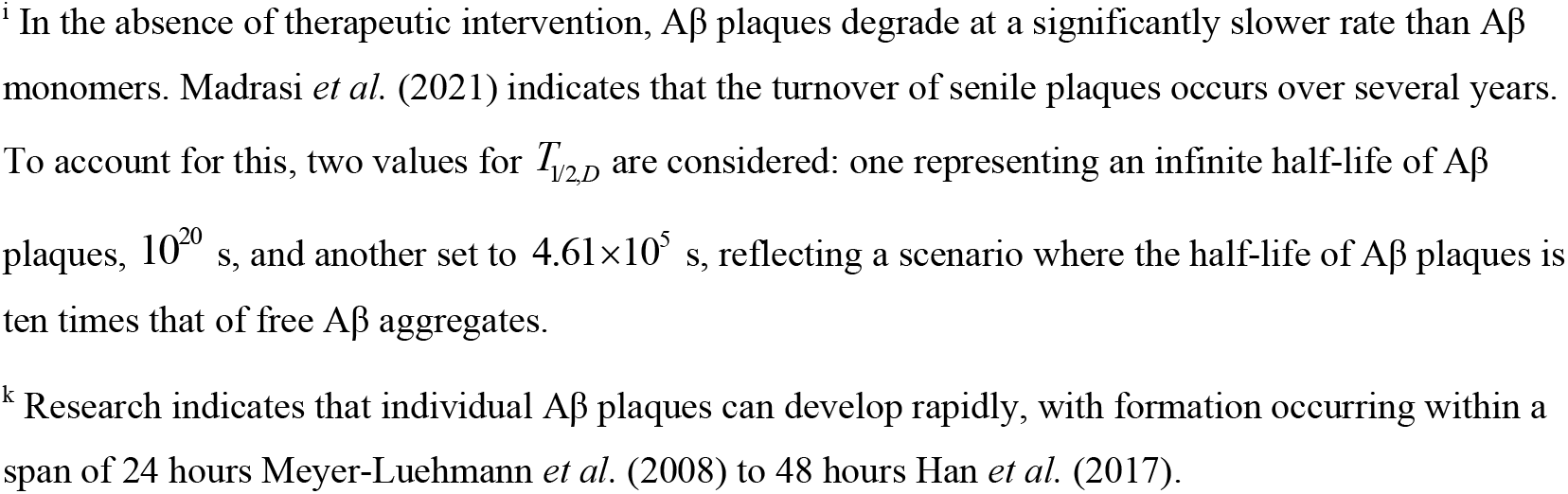
List of model parameters.

**Fig. 1.**
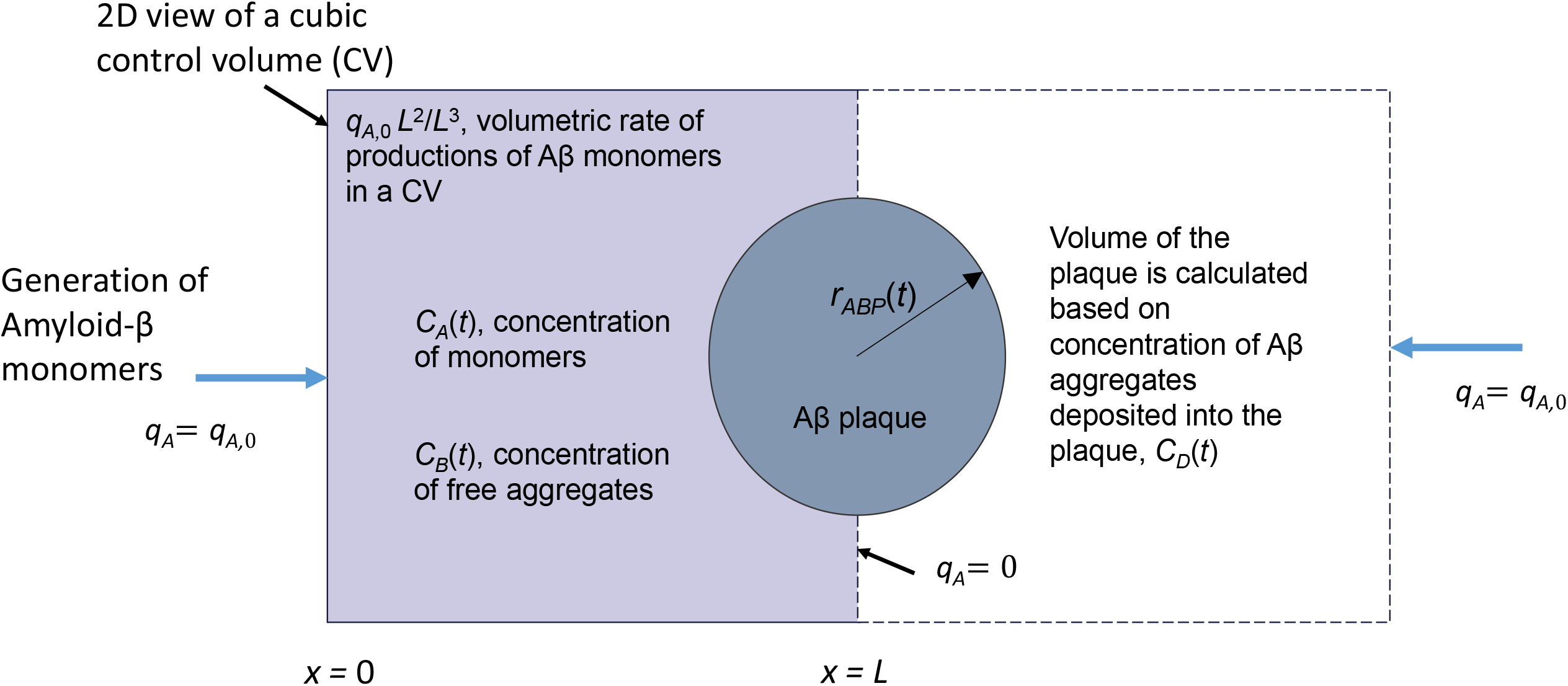
A 2D view of two cubic control volumes (CVs) with side length *L* used to model the aggregation of Aβ monomers into free aggregates and their eventual deposition into senile plaques (Liu *et al*., 2017), (Ikonomovic *et al*., 2008). Aβ monomers are assumed to be generated at lipid membranes. In the shaded CV, the production occurs at the left face (*x*=0), which has an area of *L*^2^. The right boundary of the shaded CV, *x*=*L*, is treated as symmetric, preventing Aβ monomer flux across it. A senile plaque is assumed to form at the interface between two adjacent CVs at *x*=*L*. A lumped capacitance approximation is utilized, simplifying the governing equations by assuming that all Aβ concentrations depend only on time.

By scaling governing equations, Kuznetsov (2024) introduced the following dimensionless parameter:

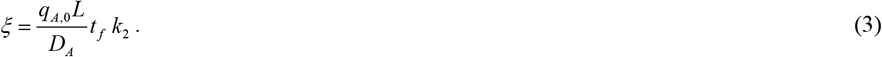

The parameter *ξ* represents the ratio of the variation in Aβ monomer concentration across the CV to the average concentration of Aβ monomers within the CV at time *t*_*f*_ . As demonstrated in Kuznetsov (2024), the lumped capacitance approximation, which assumes that Aβ concentrations depend only on time and not on location within the CV, is valid when *ξ* ≪ 1 . In Eq. (3), the diffusivity of Aβ monomers is denoted by *D*_*A*_ . Using a diffusivity value of 62.3 μm^2^/s (Waters, 2010) and the other parameter values given in Table 2, *ξ* is estimated to be 0.026. This confirms that the criterion *ξ* ≪ 1 is satisfied, and the spatial variation of Aβ monomer concentration in the CV can be neglected.

The model proposed in Kuznetsov (2025a) is adopted. By applying the conservation of Aβ monomers within the CV depicted in Fig. 1, the following equation is obtained:

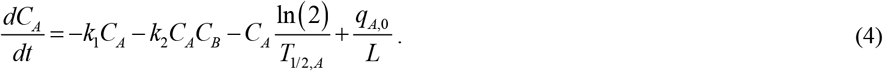

In Eq. (4), *t* represents time, which is also synonymous with calendar age. The first term on the right-hand side of Eq. (4) represents the conversion rate of Aβ monomers into aggregates driven by nucleation. The second term describes the conversion rate through autocatalytic growth. The third term accounts for the degradation of Aβ monomers, where *T*_1/2,*A*_ is an effective parameter that, in addition to intrinsic degradation, also accounts for non-intrinsic degradation mediated by microglia and macrophages. The fourth term models the production of Aβ monomers on lipid membranes. It should be noted that 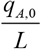 represents the average volumetric generation rate of Aβ monomers within the CV.

Applying the conservation of free Aβ aggregates within the CV yields

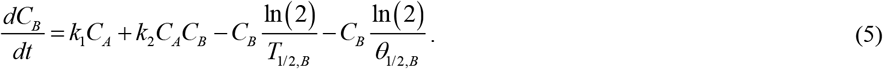

In Eq. (5), the first two terms on the right-hand side mirror the first two terms of Eq. (4) but with opposite signs. The third term represents the degradation rate of free Aβ aggregates, where *T*_1/2,*B*_ denotes an effective parameter that, in addition to intrinsic degradation, also incorporates non-intrinsic degradation mediated by microglia and macrophages. The fourth term represents the rate at which these aggregates are deposited into senile plaques.

The formation of Aβ plaques from adhesive Aβ fibrils is modeled similarly to colloidal suspension coagulation (Boltachev & Ivanov, 2020). In the model, free Aβ aggregates (*B*) are assumed to have a half-deposition time of *θ*_1/ 2, *B*_ . These aggregates deposit into Aβ plaques, and the concentration of the deposited aggregates is denoted as *C*_*D*_ . These assumptions yield the following conservation equation for Aβ aggregates incorporated into amyloid plaques:

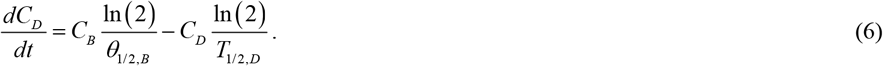

In Eq. (6), the first term on the right-hand side corresponds to the fourth term in Eq. (5) with the opposite sign, while the second term represents the half-life of Aβ aggregates within the senile plaques. Note that ^*T*^_1/ 2,*D*_ is an effective parameter that, in addition to intrinsic degradation, also accounts for non-intrinsic degradation of deposited Aβ aggregates mediated by microglia and macrophages.

The initial conditions are as follows:

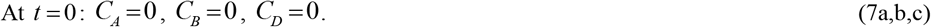

The assumption of zero initial concentration of Aβ monomers, *C*_*A*_, is an approximation. This is justified by the study’s focus on Aβ concentrations at large times, where the effect of the initial monomer concentration becomes negligible.

The time-dependent behaviors of Aβ monomers, aggregates, and plaque half-lives were modeled using two scenarios. In the first scenario, the half-lives remained constant, independent of calendar age:

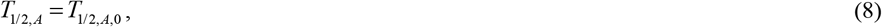

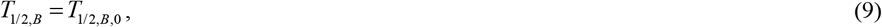

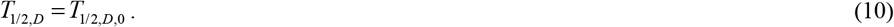

In the second scenario, the half-lives increase exponentially with calendar age, reflecting the progressive decline in proteolytic machinery function over time:

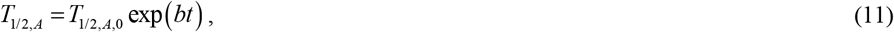

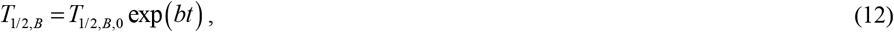

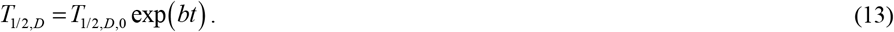

The value *b* =0.0183 hour^−1^ represents a situation in which the half-lives increase 2.5-fold over a 50-year period, consistent with the findings of Patterson *et al*. (2015).

### 2.2. Analytical solution for the scenario with *T*_1/2, *A*_ →∞, *T*_1/2,*B*_ →∞, and *T*_1/2,*D*_ →∞

For the scenario with *T*_1/2, *A*_ →∞ and *T*_1/2,*B*_ →∞, the numerical solutions presented in Figs. 2 and 3 indicate that the concentrations of Aβ monomers and oligomers, *C*_*A*_ and *C*_*B*_, respectively, approach asymptotic steady-state values over time. This section determines these asymptotic values. The solution derived here is labeled as “analytical” in the figures. The steady-state values of *C*_*A*_ and *C*_*B*_ are obtained from the following equations, derived from applying these conditions to Eqs. (4) and (5):

**Fig. 2.**
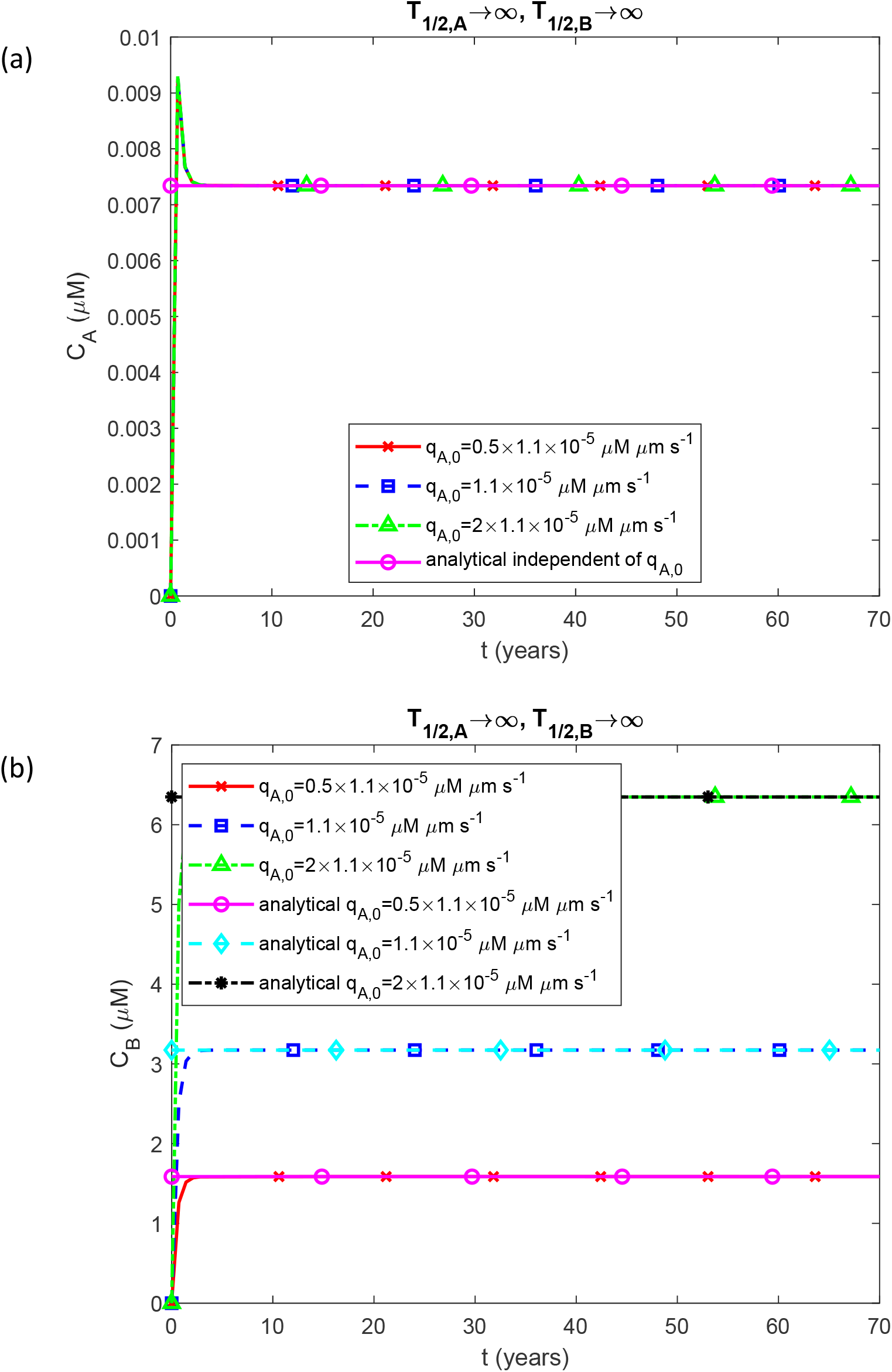
Comparison of numerical and analytical results for the molar concentration of (a) Aβ monomers, *C*_*A*_, and (b) Aβ oligomers, *C*_*B*_, as a function of time for different values of Aβ monomer flux into the CV, *q*_*A*,0_ . Parameters: *T*_1/2, *A*_ =10^20^ s, *T*_1/2, *B*_ =10^20^ s. The plotted analytical solutions are derived from Eqs. (17) and (18). Note that in each plotted case *C*_*A*_ and *C*_*B*_ reach constant asymptotic values as time increases. This is essential for deriving the analytical solution.

**Fig. 3.**
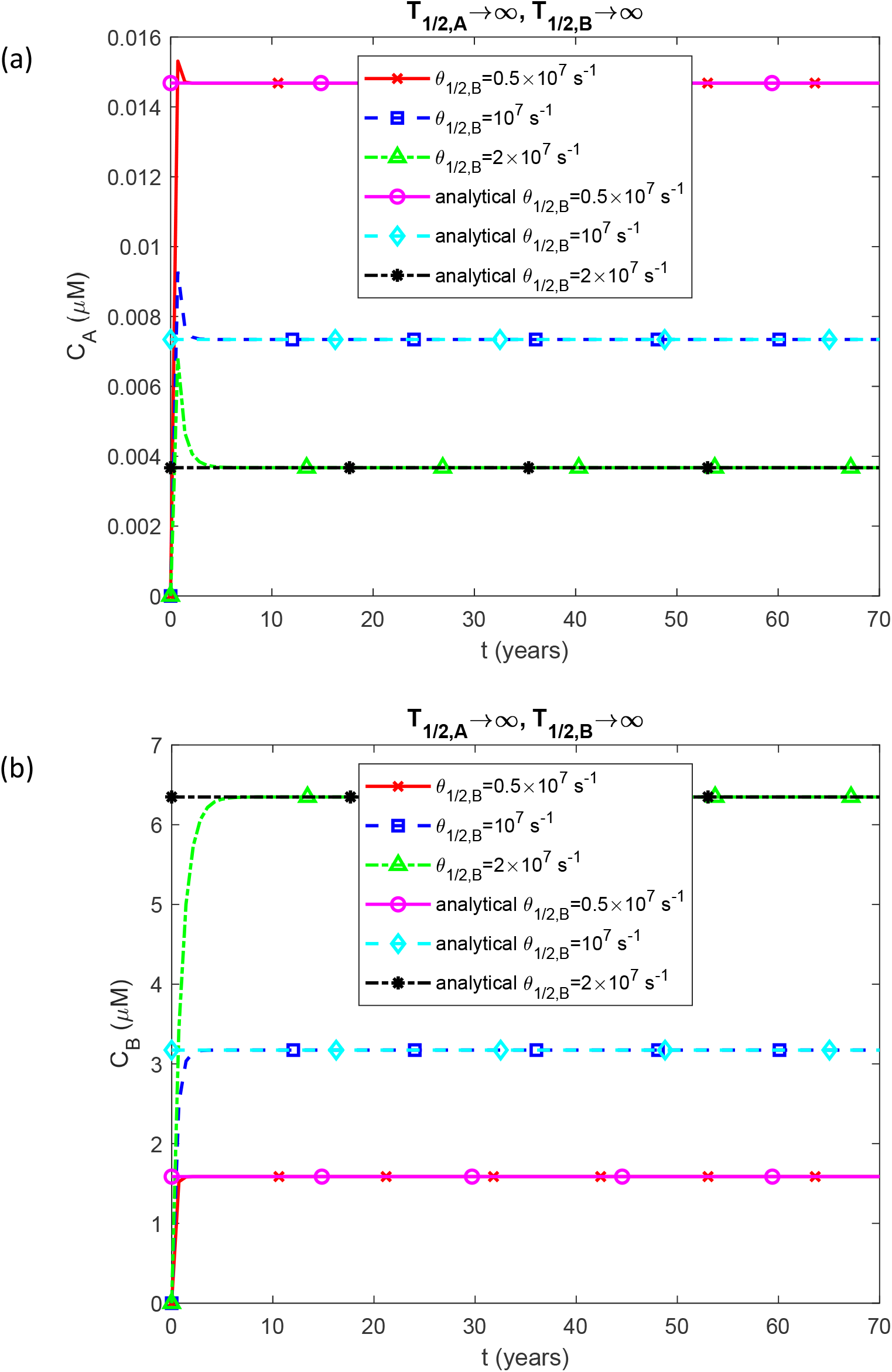
Comparison of numerical and analytical results for the molar concentration of (a) Aβ monomers, *C*_*A*_, and (b) Aβ oligomers, *C*_*B*_, as a function of time for different values of half-deposition times of free Aβ aggregates into senile plaques, *θ*_1/2,*A*_ . Parameters: *T*_1/2,*A*_ =10^20^ s, *T*_1/2,*B*_ =10^20^ s. The plotted analytical solutions are derived from Eqs. (17) and (18). Note that in each plotted case *C*_*A*_ and *C*_*B*_ reach constant asymptotic values as time increases. This is essential for deriving the analytical solution.

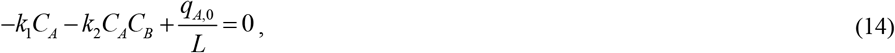

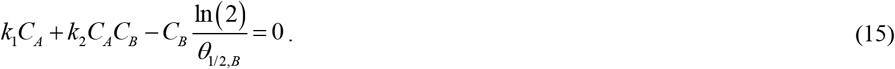

Adding Eqs. (14) and (15) yields:

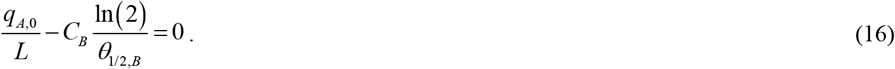

Solving Eq. (16) for *C*_*B*_ results in:

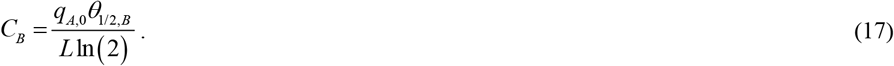

Substituting Eq. (17) into Eq. (14) and solving for *C*_*A*_ yields:

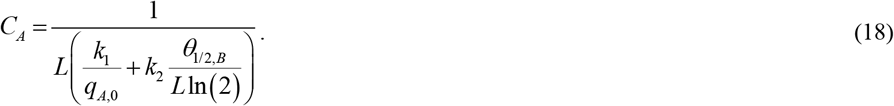

In agreement with Eq. (17), Figs. 2b and 3b demonstrate that the asymptotic value of *C*_*B*_ obtained for *t* →∞ is affected by both *q*_*A*,0_ and *θ*_1/ 2, *B*_ . While the asymptotic value of *C*_*A*_ remains unaffected by *q*_*A*,0_ for small values of *k*_1_, it does depend on the value of *θ*_1/ 2, *B*_ (Figs. 2a and 3a). This result is consistent with Eq. (18). The numerical and analytical solutions are in excellent agreement when time exceeds five years (Figs. 2 and 3).

For *T*_1/ 2,*D*_ →∞, Eq. (6) can be rewritten as:

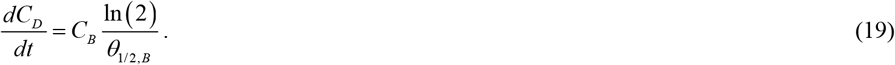

Substituting Eq. (17) into Eq. (19) yields:

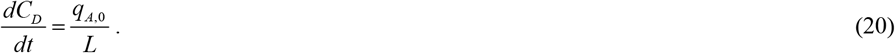

Integrating Eq. (20) with the initial condition (7c) gives:

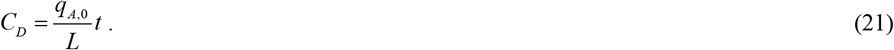

According to Eq. (21), *C*_*D*_ increases linearly with time, depending on *q*_*A*,0_ (Fig. S1a) while remaining independent of *θ*_1/ 2, *B*_ S2a). The numerical and analytical solutions are nearly identical (Figs. S1a and (Fig. S2a).

An equation for the radius of senile plaques as a function of *C*_*D*_ is derived in Section S1 of the Supplemental Materials, using the approach developed in Watzky *et al*. (2008). Substituting Eq. (21) into Eq. (S5) gives an analytical solution for the scenario with *T*_1/ 2, *A*_ →∞, *T*_1/ 2,*B*_ →∞, and *T*_1/ 2,*D*_ →∞ :

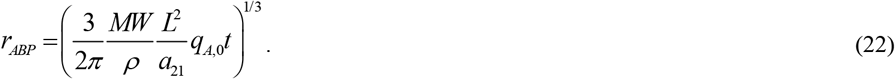

According to Eq. (22), the plaque radius grows in direct proportion to the cube root of time. It depends on *q*_*A*,0_ (Fig. S1b) but is independent of *θ*_1/ 2, *B*_ (Fig. S1b). The numerical and analytical solutions for the plaque radius are in excellent agreement (Figs. S1b and S2b).

Unlike numerical simulations, the analytical solutions explicitly reveal scaling laws: monomer and oligomer concentrations saturate at steady-state values for large times, while plaque radius grows as the cube root of time ( *t*^1/3^). These relationships emerge directly from the analytical solutions but would require additional parametric analysis to extract from numerical simulations.

### 2.2. Analytical solution for the scenario with constant finite half-lives

*T*_1/ 2, *A*_, *T*_1/ 2,*B*_, and *T*_1/2,*D*_

The numerical solutions of Eqs. (4) and (5) with initial conditions (7a, b) for finite, time-independent values of *T*_1/ 2, *A*_ and *T*_1/ 2,*B*_, as shown in Figs. S3 and S4, indicate that *C*_*A*_ and *C*_*B*_ reach steady-state concentrations over time. These asymptotic concentrations of *C*_*A*_ and *C*_*B*_ can be determined by solving the following steady-state equations, derived from Eqs. (4) and (5):

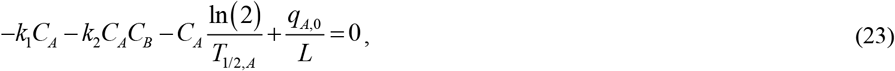

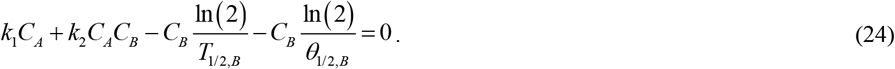

The analytical solution for this case, which is labeled as “full analytical” in the figures, is derived in Section S2 of the Supplemental Materials.

### 2.2. Criterion of accumulated neurotoxicity of Aβ oligomers and proposed biomarker for biological aging

The precise mechanism underlying the neurotoxicity of Aβ oligomers remains unclear. Soluble Aβ oligomers exert neurotoxic effects through multiple mechanisms, including direct interactions with neuronal membranes and synaptic receptors, which lead to disruption of membrane integrity, ionic homeostasis, and receptor function, as well as the activation of inflammatory pathways. They also induce oxidative stress by forming metal–Aβ complexes, trigger mitochondrial dysfunction, and cause excessive calcium influx, ultimately impairing neuronal viability. Furthermore, their ability to propagate between cells and accumulate in subcellular compartments contributes to the spread of neuropathology and progressive synaptic failure in AD (Zhao *et al*., 2012; Sengupta *et al*., 2016; Lee *et al*., 2017; Chen *et al*., 2017; Mrdenovic *et al*., 2022; Benilova *et al*., 2012).

Certain Aβ oligomers, such as Aβ42, may possess inherently greater neurotoxicity (Mrdenovic *et al*., 2022; Gu & Guo, 2013), whereas in other cases, neurotoxic effects may arise from a heterogeneous mixture of oligomers differing in structure, stability, and concentration. If neurotoxicity is nonspecific and results from the combined effects of various oligomers, a parameter quantifying the accumulated time-dependent damage caused by Aβ oligomers can be defined, following Kuznetsov (2025a), Kuznetsov (2025c), as:

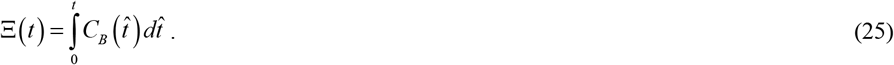

It should be noted that the governing equations are formulated based on mass balance within a CV shown in Fig. 1, which has a characteristic dimension *L* ≈ 50 μm (half the inter-plaque spacing). Consequently, the accumulated neurotoxicity represents neural damage at the local (cellular) level rather than a whole-brain average.

Because the model parameters—specifically the production flux ( *q*_*A*,0_) and the proteolytic half-lives ( *T*_1/2,*A*,0_ and *T*_1/2,*B*,0_)—can be tuned to reflect different physiological conditions, the developed mathematical framework is capable of representing distinct brain regions. For instance, regions that remain “spared” in early AD would be characterized by highly efficient protein degradation (short half-lives), whereas “vulnerable” regions such as the hippocampus or entorhinal cortex would be modeled using longer half-lives or elevated production rates. This mathematical framework thus provides a mechanistic tool for exploring why certain brain regions exhibit more severe AD pathology and neurodegeneration than others at the same chronological age.

By substituting the analytical solution for *C*_*B*_ from Eq. (17), which holds for *T*_1/ 2, *A*_ →∞ and *T*_1/ 2,*B*_ →∞, into Eq. (25), the following expression is derived:

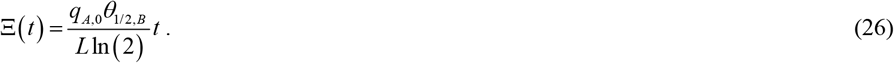

The biological age of neurons (hereafter referred to as biological age) is defined by:

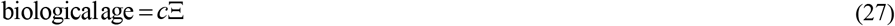

Substituting the analytical solution from Eq. (26) into Eq. (27) results in:

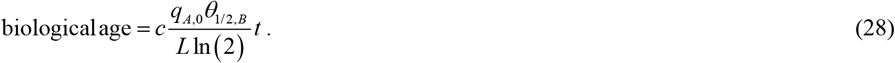

An interesting consequence of Eq. (28) is that biological age exhibits a linear dependence on the half-deposition time of free Aβ aggregates into senile plaques, *θ*_1/2,*B*_ .

While a few Aβ-clearing drugs, such as aducanumab and lecanemab, have received FDA approval, numerous clinical trials of similar therapies for AD treatment have failed (Zhang *et al*., 2023; Xiao & Zhang, 2024). Even approved agents substantially reduce plaque burden yet produce only modest clinical benefits without reversing established cognitive impairment in symptomatic patients (Musiek *et al*., 2021).

The principal advantage of defining biological age as directly proportional to accumulated neurotoxicity (Eq. (27)) is that this formulation inherently encodes irreversibility, providing a theoretical explanation for this therapeutic gap. Because biological age is defined as a time integral (Eqs. (25) and (27)), therapeutic removal of plaques may reduce current oligomer concentration but cannot “undo” accumulated damage. Even if oligomer levels were reduced to zero, biological age would continue from its current value rather than resetting. The integral structure mathematically proves that biological age can only increase or plateau—never decrease.

This framework suggests that observable plaque pathology is merely a secondary indicator, whereas the primary driver of neuronal aging is cumulative historical exposure to toxic oligomers. The model thus explains why plaque clearance does not restore cognitive function: the neuronal biological clock has already advanced irreversibly through decades of oligomer exposure. Therapeutic interventions can slow the rate at which biological age increases but cannot reverse damage already integrated.

Another advantage of the proposed definition is that it renders neuronal biological age path-dependent rather than merely state-dependent. Consider two neurons that reach identical final biomarker profiles at the same calendar age. Despite this similarity in observable pathology, these neurons may have different biological ages if their accumulated neurotoxicity differs—for instance, one neuron may have experienced high oligomer concentrations over a short duration, while the other sustained lower concentrations over an extended period.

This path-dependence provides a theoretical explanation for clinical heterogeneity: individuals with similar biomarker measurements can exhibit markedly different cognitive outcomes because their historical exposure trajectories differ. Longitudinal amyloid-PET studies demonstrate substantial heterogeneity in the rate and timing of Aβ accumulation, revealing that individuals with similarly positive amyloid-PET scans can have vastly different risks and rates of future cognitive decline depending on how rapidly their pathology developed (Bollack *et al*., 2024). This temporal dimension—invisible in single-timepoint assessments—is explicitly encoded in the integral biomarker, which captures the cumulative exposure history that determines true biological age.

Note that the analytical solution for oligomer concentration (Eq. (17)) predicts that *C*_*B*_ remains approximately constant throughout most of the lifespan after an initial transient period. The numerical solutions displayed in Figs. 2b, 3b, and 5b confirm this behavior: *C*_*B*_ starts at zero at birth but rapidly reaches a steady-state value within the first few years. Consequently, calculating accumulated neurotoxicity would require only a single measurement of oligomer concentration combined with knowledge of the individual’s calendar age, since the 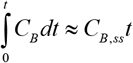 for most of the lifespan. Methods for measuring oligomer concentration in human brain tissue already exist (Yang *et al*., 2013), though these currently require post-mortem or biopsy samples.

### 2.3. Analysis of the sensitivity of biological age to various parameters

To assess the sensitivity of biological age to model parameters, local sensitivity coefficients were calculated. These coefficients represent the first-order partial derivatives of biological age with respect to the model parameters (Beck & Arnold, 1977; Zadeh & Montas, 2010; Zi, 2011; Kuznetsov & Kuznetsov, 2019). The dimensionless relative sensitivity coefficients were determined following the methodology outlined in (Zadeh & Montas, 2010; Kacser *et al*., 1995). For instance, the dimensionless sensitivity of biological age to the flux of Aβ monomers into the CV is given by:

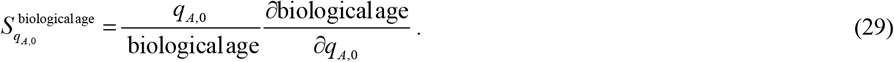

Utilizing the analytical solution given by Eq. (28) yields the following expression for the sensitivity of biological age to calendar age *t*:

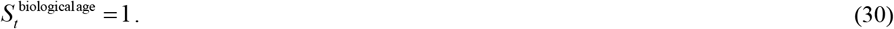

Eq. (30) is consistent with the fact that biological age increases in direct proportion to calendar age *t*; see Eq. (28). Also,

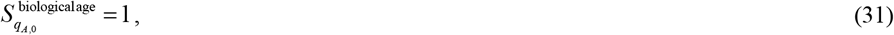

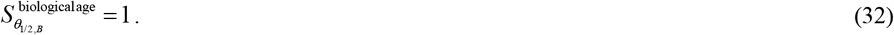

The parameter 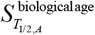 is calculated using the same parameter values as those used for one of the curves in Fig. 4a: *T*_1/2, *A*_ = 4.61×10^3^ s (the biologically relevant value) and *T*_1/2, *B*_ =10^20^ s. Utilizing Eq. (S16) yields:

**Fig. 4.**
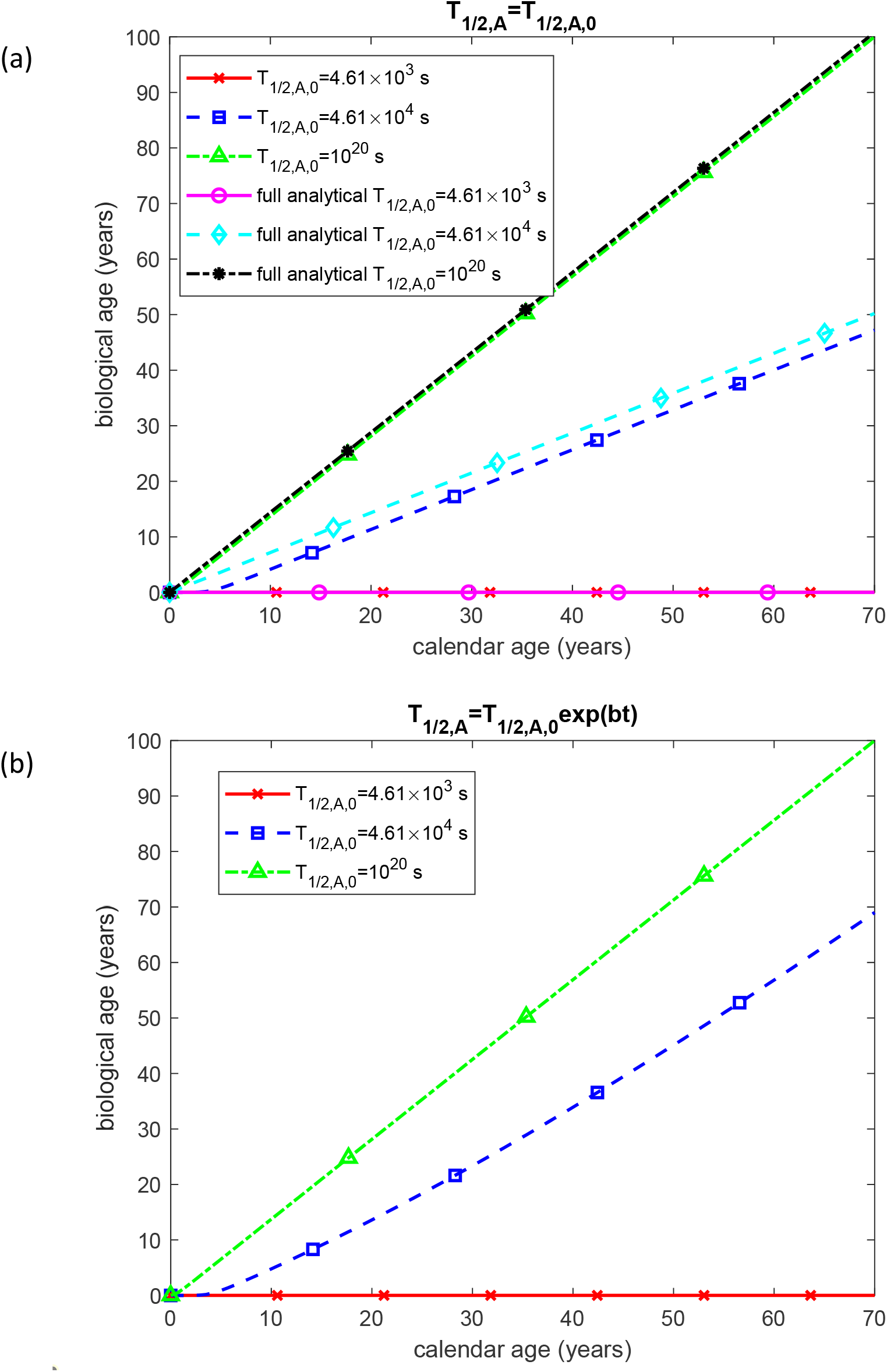
Biological age as a function of calendar age for different half-lives of Aβ monomers, *T*_1/2,*A*_. (a) The half-life of Aβ monomers, *T*_1/2,*A*_, remains constant and independent of calendar age. (b) The half-life of Aβ monomers, *T*_1/2,*A*_, increases exponentially with calendar age. Parameters: *T*_1/2,*B*_ =10^20^ s.

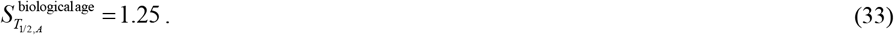

Similarly, 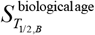 is calculated using the same parameter values as those used for one of the curves in Fig. 6a: *T*_1/2, *A*_ =10^20^ s and *T*_1/2, *B*_ = 4.61×10^4^ s (the biologically relevant value). Using Eq. (S17) yields:

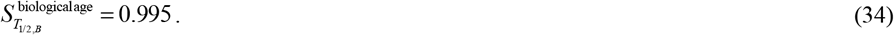

## 3. Results

The details of the numerical solution are outlined in Section S3 of the Supplementary Materials. Unless otherwise specified in a figure or its caption, parameter values are taken from Table 2. Two values are employed in the computations for the half-lives of monomers, oligomers, and plaques (*T*_1/2,*A*,0_, *T*_1/2,*B*,0_, and *T*_1/2,*D*,0_): a physiologically relevant value and an infinitely large value, the latter representing the assumption of complete failure of the degradation machinery. In the numerical implementation, the infinitely large value is approximated as 10^20^ s.

Accumulated neurotoxicity, Ξ, quantifies the neuronal damage caused by Aβ oligomers. Therefore, it is proposed as a biomarker for neuronal biological age. Biological age is assumed to be directly proportional to accumulated neurotoxicity. The maximum accumulated neurotoxicity value, 6.95×10^9^ µMꞏs in Fig. S5a, is assumed equivalent to a biological age of 100 years, as defined by Eq. (27).

If the proteolytic machinery responsible for Aβ monomer degradation remains functional ( *T*_1/2, *A*_ = 4.61×10^3^ s, a physiologically relevant value), neuronal aging due to Aβ oligomer neurotoxicity remains minimal (Fig. 4a). This does not imply that neurons will not age, but rather that Aβ oligomer neurotoxicity will not be a significant contributing factor in aging. In contrast, if Aβ monomer degradation is entirely impaired ( *T*_1/2, *A*_ =10^20^ s), the resulting neurotoxicity will drive neurons to a biological age of 100 years by a calendar age of 70 years. If the half-life of Aβ monomers is increased tenfold beyond the physiologically relevant value ( *T*_1/2, *A*_ = 4.61×10^4^ s), neurons will reach a biological age of approximately 47 years by the time the calendar age reaches 70 years (Fig. 4a). The observed increase in biological age with the increasing half-life of Aβ monomers (Fig. 4a) is consistent with a positive value of 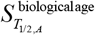; see Eq. (33).

For *T*_1/2, *A*_ = 4.61×10^3^ s and *T*_1/2, *A*_ =10^20^ s, the numerical and full analytical solutions for the biological age (given by Eq. (S15)) are in excellent agreement. However, for *T*_1/2, *A*_ = 4.61×10^4^ s, a systematic discrepancy between the two solutions persists over time (Fig. 4a). To understand the origin of this discrepancy, Fig. 5 plots the concentration of Aβ oligomers as a function of time for the scenario depicted in Fig. 4a. As seen in Fig. 5, the discrepancy arises from the difference between the numerical and full analytical solutions (given by Eq. (S10)) for *C*_*B*_ during the first 10 years. Although the agreement between the two solutions improves after this period, the initial discrepancy leads to a persistent offset in the accumulated neurotoxicity (Fig. S5a). This disparity occurs because accumulated neurotoxicity is computed as the integral of the oligomer concentration over time (Eq. (25)), and during the first 10 years, the numerically predicted oligomer concentration is lower than the analytically predicted one (Fig. 5).

**Fig. 5.**
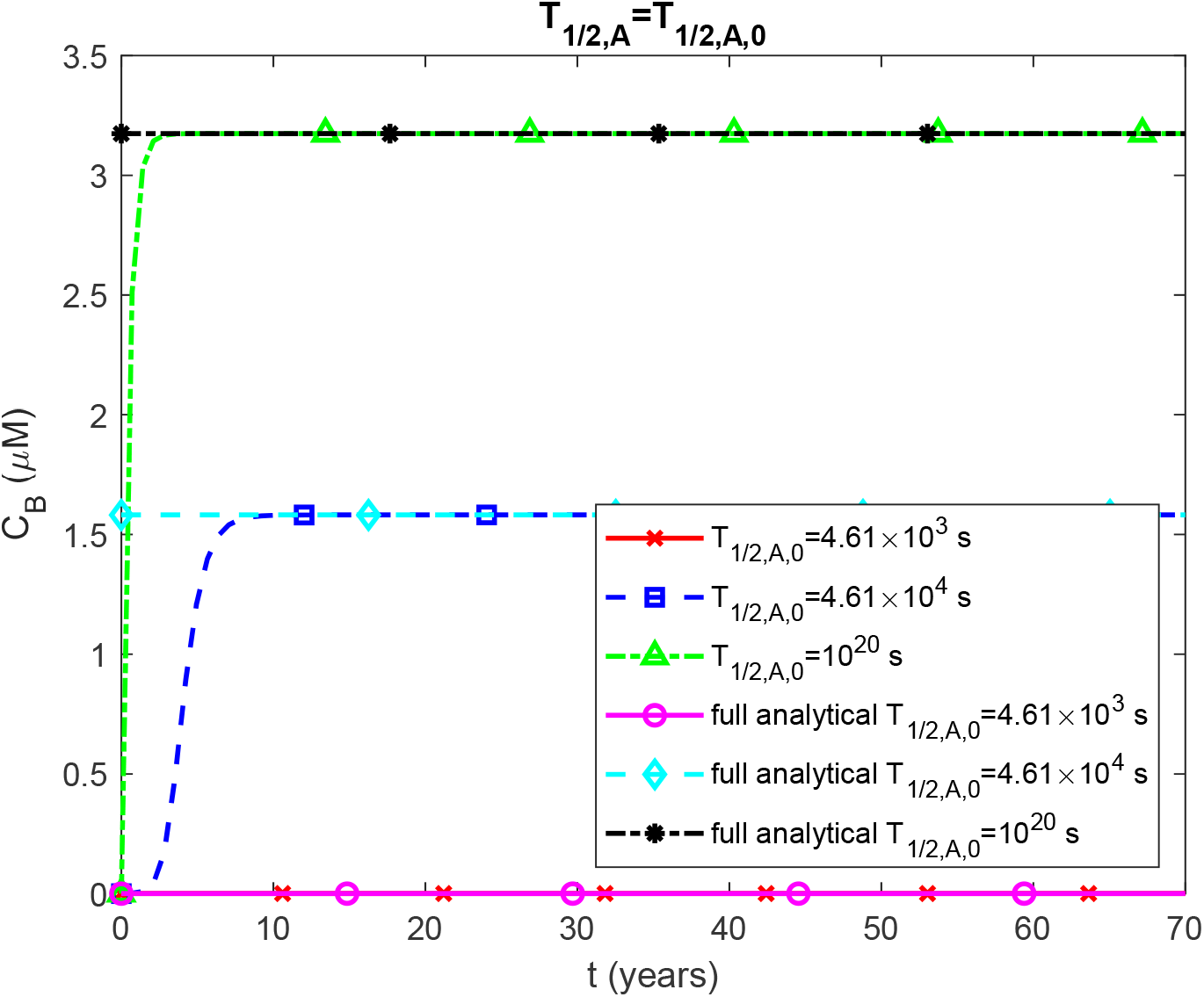
Molar concentration of Aβ oligomers, *C*_*B*_, as a function of time for different half-lives of Aβ monomers, *T*_1/2,*A*_. This figure corresponds to the scenario shown in Fig. 4 and uses the same parameter values. The full analytical solution, obtained from Eq. (S10), is plotted. Notably, a significant discrepancy between the numerical and full analytical solutions is observed during the first 10 years for the case where *T*_1/2,*A*,0_ = 4.61×10^4^ s. However, in all cases, *C* eventually reaches a constant asymptotic value over time. Parameters: *T*_1/2,*A*_ =10^20^ s.

**Fig. 6.**
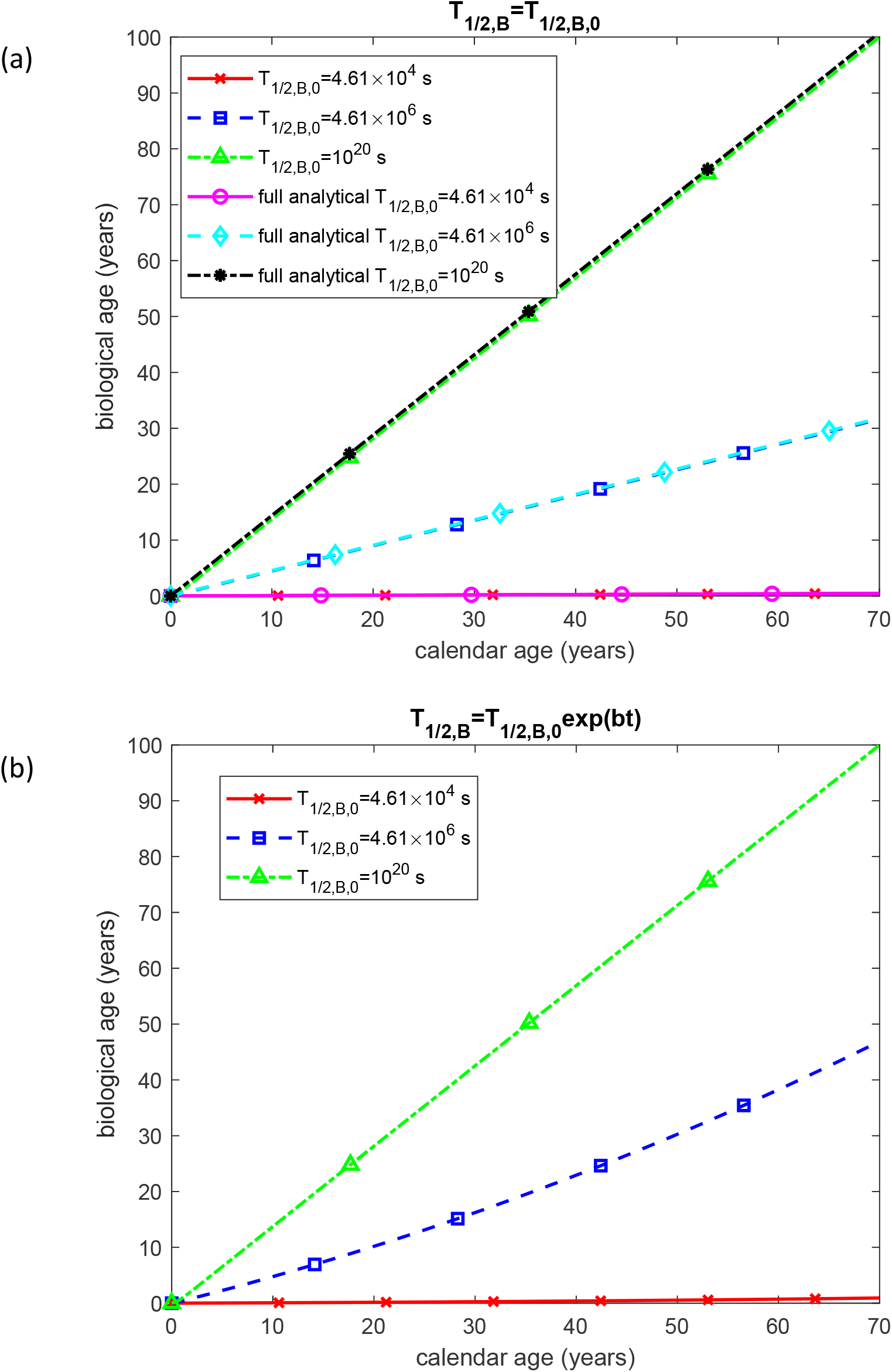
Biological age as a function of calendar age. (a) The half-life of free Aβ aggregates, *T*_1/2,*B*_, remains constant over time. (b) The half-life of free Aβ aggregates, *T*_1/2,*B*_, increases exponentially with calendar age. Parameters: *T*_1/2,*A*_ =10^20^ s.

Fig. 4b depicts the scenario in which the half-life of Aβ monomers, *T*_1/2, *A*_, increases exponentially with calendar age from its initial value, *T*_1/2, *A*,0_, at *t*=0. The exponential growth of *T*_1/2, *A*_ / *T*_1/ 2, *A*,0_, represented by exp(*bt*), is shown in Fig. S6, where the parameter *b* is estimated based on data from (Patterson *et al*., 2015). This model is plausible, as the Aβ monomer degradation machinery is expected to function efficiently in youth but gradually decline with age, leading to an age-dependent increase in the half-life of Aβ monomers. Under this scenario, if *T*_1/2, *A*,0_ is set to 4.61×10^4^ s, the biological age at a calendar age of 70 years reaches approximately 70 years (Fig. 4b), compared to about 47 years for the same *T*_1/2, *A*,0_ without exponential growth over time (Fig. 4a).

When the half-life of Aβ oligomers is at a physiologically relevant value of 4.61×10^4^ s, the degradation machinery functions efficiently, resulting in a very insignificant increase in biological age with calendar age (Fig. 6a). However, increasing the half-life to 4.61×10^6^ s accelerates aging, raising the biological age to approximately 32 years by the time the calendar age reaches 70 years. If the half-life becomes infinitely large, the biological age reaches 100 years at a calendar age of 70 years (Fig. 6a). The increase of the biological age with *T*_1/2, *B*_ observed in Fig. 6a agrees with the positive sensitivity 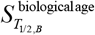; see Eq. (34).

If the half-life of Aβ oligomers increases exponentially with age, an initial value of 4.61×10^6^ s results in a biological age of approximately 47 years at a calendar age of 70 years (Fig. 6b), compared to 32 years for the same value of *T*_1/2,*B*,0_ in Fig. 6a. The trends in accumulated neurotoxicity (Fig. S7) closely follow those in Fig. 6 for biological age, as the model assumes a direct proportionality between biological age and accumulated neurotoxicity; see Eq. (27).

Variations in the half-life of Aβ aggregates within senile plaques do not affect biological age (Fig. S8a,b). This is because the model assumes that accumulated neurotoxicity is driven by Aβ oligomers, whereas senile plaques, considered non-toxic, serve as relatively benign reservoirs for these oligomers—a hypothesis consistent with (Gouras *et al*., 2015; Cline *et al*., 2018).

Fig. 7a illustrates the effect of Aβ monomer flux into the CV, *q*_*A*,0_, representing the production rate of Aβ monomers, on biological age. Aβ monomers are generated at lipid membranes, and an increase in their flux significantly accelerates biological aging. When the flux is reduced to half of its physiologically relevant value ( 0.5 ×1.1×10^−5^ μM μm s^-1^), the biological age at a calendar age of 70 years is approximately 50 years (Fig. 7a). At a physiologically relevant value of the flux (1.1×10^−5^ μM μm s^-1^), the biological age at 70 years reaches 100 years. If the flux is doubled from its physiologically relevant value ( 2 ×1.1×10^−5^ μM μm s^-1^), the biological age reaches 100 years when calendar age is only 35 years (Fig. 7a).

**Fig. 7.**
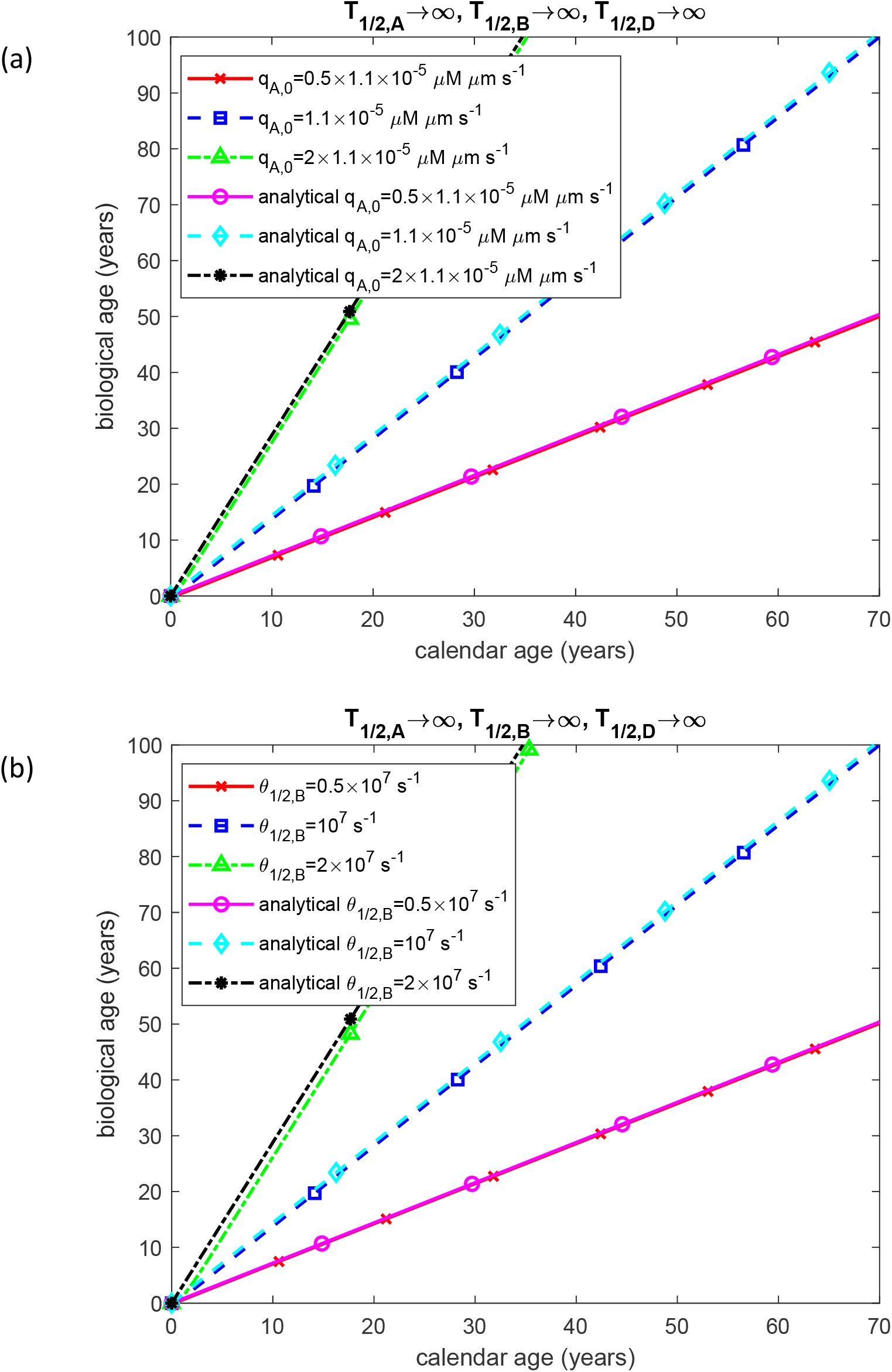
(a) Biological age as a function of calendar age for different values of the flux, *q*_*A*,0_, of Aβ monomers per unit area at the left-hand boundary of the CV shown in Fig. 1. (b) Biological age as a function of calendar age for different values of the half-deposition time, *θ*_1/2,*B*_, of free Aβ aggregates into senile plaques. Parameters: *T*_1/2,*A*_ =10^20^ s, *T*_1/2,*B*_ =10^20^ s.

Fig. 7b illustrates the effect of the half-deposition time of free Aβ aggregates into senile plaques, *θ*_1/2,*B*_, on biological age. Interestingly, its effect is analogous to that of the Aβ monomer flux: when *θ*_1/2,*B*_ is reduced to half of its physiologically relevant value, the biological age at a calendar age of 70 years is approximately 50 years (Fig. 7b). At the physiologically relevant value of *θ*_1/2,*B*_, the biological age at 70 years reaches 100 years. When *θ*_1/2,*B*_ is doubled from its physiologically relevant value, the biological age reaches 100 years as early as a calendar age of 35 years (Fig. 7b).

Figs. 7a and 7b demonstrate strong agreement between the numerical and analytical solutions; hence, the analytical solution given by Eq. (28) can be used to explain this similarity. Eq. (28) shows that biological age is proportional to the product of *q*_*A*,0_ and *θ*_1/ 2, *B*_ rather than these parameters independently. This explains the striking similarity between Figs. 7a and 7b—halving monomer production has the identical effect as halving the half-deposition time. This equivalence was completely non-obvious from numerical simulations and has important therapeutic implications: interventions targeting either Aβ production or plaque deposition kinetics will have equivalent effects on biological aging. Consequently, variations in these parameters produce identical effects on biological age. The effects of *q*_*A*,0_ and *θ*_1/ 2, *B*_ on accumulated neurotoxicity follow the same trends, as biological age is assumed to be directly proportional to accumulated neurotoxicity (Figs. S9a,b).

Biological age exhibits minimal dependence on the nucleation rate constant, *k*_1_ (Fig. 8a). However, it increases with an increase in the autocatalytic growth rate constant, *k*_2_, indicating that Aβ oligomer production is primarily driven by the autocatalytic process (Fig. 8b). For biologically relevant values of *k*_2_ (represented by the line with red crosses), the relationship between biological age and calendar age is initially nonlinear for the first 30 years. Beyond this period, it transitions into a linear trend, consistent with Eq. (28). The curves depicting accumulated neurotoxicity vs time for different values of *k*_1_ and *k*_2_ exhibit similar trends (Fig. S10a and S10b).

**Fig. 8.**
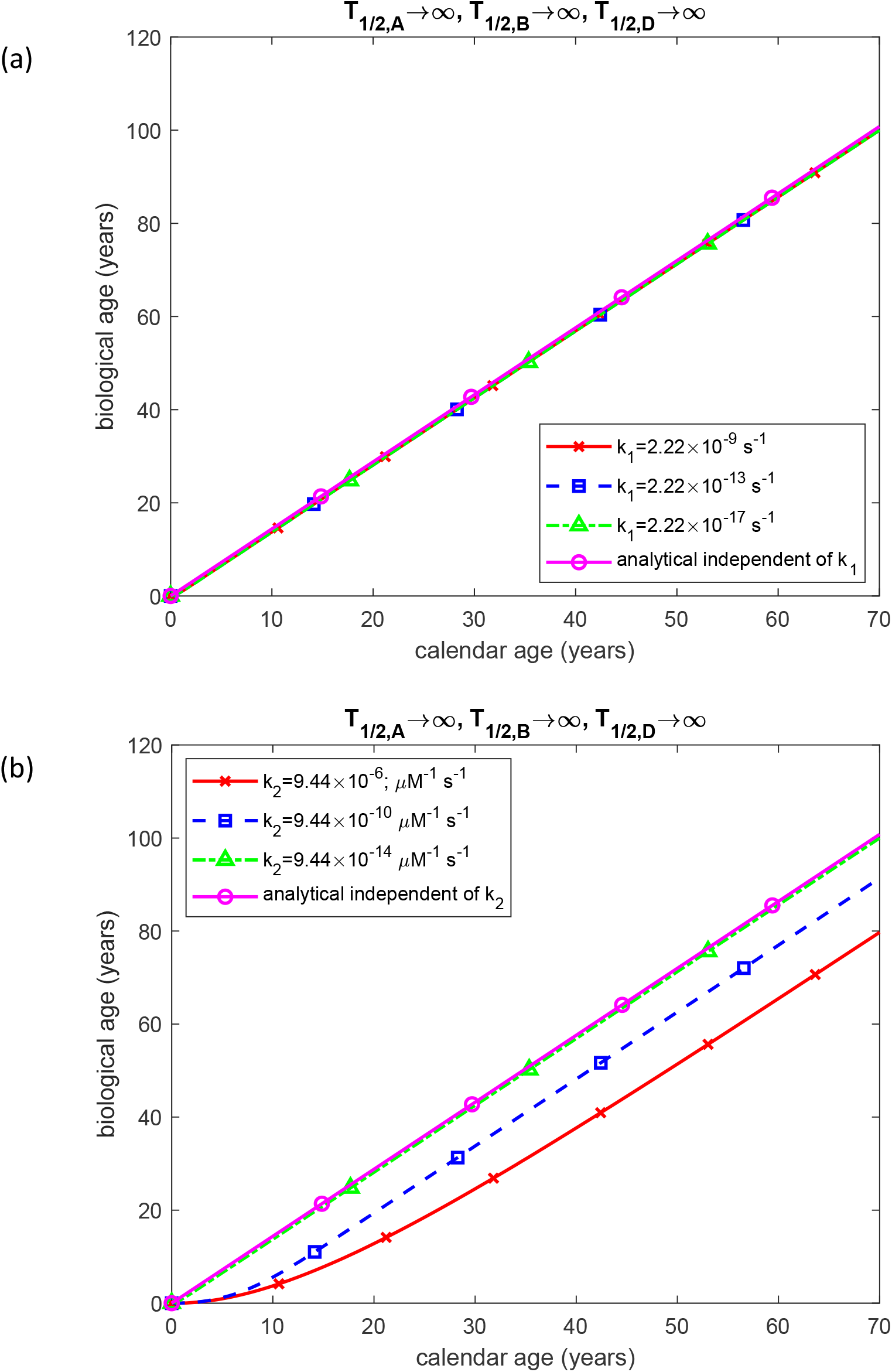
(a) Biological age as a function of calendar age for different values of the nucleation rate constant, *k*_1_, in the first pseudo-elementary reaction step of the F-W model for Aβ peptide. Parameters: *K*_2_ = 9.44 × 10^−6^ μM^-1^ s^-1^, *T*_1/2,*A*_ =10^20^ s, *T*_1/2,*B*_ =10^20^ s, *T*_1/2,*D*_ =10^20^ s. (b) Biological age as a function of calendar age for different values of the autocatalytic growth rate constant, *k*_2_, in the second pseudo-elementary reaction step of the F-W model for Aβ peptide. Parameters: *k*_1_ = 2.22×10^-9^ s^-1^, *T*_1/2,*A*_ =10^20^ s, *T*_1/2,*B*_ =10^20^ s, *T*_1/2,*D*_ =10^20^ s.

## 4. Discussion, limitations of the model, and future directions

A central question addressed in this study is the definition of aging and how healthy aging differs from pathology. This paper adopts a practical framework in which aging is understood as the progressive loss of the organism’s ability to recover from multisystem damage accumulated with increasing calendar age. Cataract formation, for instance, exemplifies such age-related damage.

Within this framework, a criterion was introduced to characterize the accumulated neurotoxicity of Aβ oligomers in the AD brain. This criterion serves as a potential biomarker of neuronal aging and may prove valuable for assessing the efficacy of various anti-aging therapeutic interventions. However, its limitation lies in the potential inability to discriminate between, for example, neurons in a 70-year-old individual with AD and those in an 85-year-old individual without neurological impairment.

Defining neuronal biological age in terms of accumulated neurotoxicity induced by Aβ oligomers provides a theoretical foundation for understanding the irreversibility of neuronal damage in AD. The proposed biomarker framework posits that neurodegeneration is driven by cumulative exposure to oligomers over a lifetime. This hypothesis explains the limited efficacy of Aβ-clearing drugs: while these therapies may reduce the current concentration of toxic species, they cannot reverse the accumulated neurotoxicity that defines a neuron’s biological age.

The model also demonstrates that neuronal biological age is path-dependent: two neurons reaching the same final plaque burden via different trajectories will have different biological ages. This path-dependence explains clinical heterogeneity (Goyal *et al*., 2018)—why individuals with similar biomarker values exhibit divergent cognitive trajectories.

Furthermore, the model illuminates why therapeutic windows exist (Nakashima *et al*., 2025). If proteolytic half-lives increase exponentially with age (Eqs. (11)-(13)), reflecting gradual failure of protein quality control machinery, biological age increases nonlinearly over time (Figs. 4b, 6b). This produces an increasingly rapid accumulation of neurotoxicity at later calendar ages. Since therapeutic interventions cannot reverse damage already accumulated, late-stage interventions are predicted to be markedly less effective than early preventive treatments, even if both achieve identical reductions in current oligomer concentrations.

The half-lives of Aβ monomers, oligomers, and aggregates serve as indicators of the efficiency of the Aβ degradation machinery. When these half-lives are assumed to be constant, biological age exhibits a linear dependence on calendar age (Figs. 4a, 6a, 7a, and 7b). This aligns with the analytical solution for biological age given by Eq. (28), which predicts a linear relationship with calendar age *t*. An exception is observed in Fig. 8b, which depicts biological age as a function of time for different values of the autocatalytic constant *k*_2_ . For biologically relevant values of *k*_2_, the curve exhibits a parabolic shape at small *t* but transitions to a linear trend at larger times. This remains consistent with Eq. (28), as the analytical solution describes the asymptotic regime at large times, where the relationship becomes linear.

If the half-lives of monomers and oligomers are assumed to increase exponentially with age, simulating the decline of proteolytic machinery over time, the relationship between biological age and calendar age becomes nonlinear for intermediate initial half-life values (Figs. 4b and 6b).

Numerical analysis revealed a similarity between the effects of Aβ monomer production rate (represented by Aβ monomer flux into the CV, *q*_*A*,0_) and the half-deposition time of free Aβ aggregates into senile plaques, *θ*_1/2,*B*_ (Fig. 7a and 7b). This similarity is explained by the analytical solution for biological age, which depends on the product *q*_*A*,0_*θ*_1/ 2, *B*_ .

This study is based on the assumption that Aβ oligomers represent the most neurotoxic species in AD. While this is a widely supported hypothesis, alternative theories regarding the causative mechanisms of AD also exist. A potential limitation of the proposed biomarker is the assumption that accumulated neurotoxicity begins to rise immediately after birth, whereas it is more likely to increase only after a certain age. Future research should focus on experimental evaluation of the proposed biomarker, which represents accumulated neurotoxicity, for assessing neuronal aging. Additionally, the potential applicability of accumulated toxicity (defined similarly) as a biomarker for aging in non-neural tissues should be explored. The influence of sex—particularly the greater Aβ deposition observed in specific brain regions among women—and the impact of the apolipoprotein E (*APOE*) ε2, ε3, and ε4 genotype on biological age warrant further investigation. Simulations incorporating an extended lifespan (80–90 years) should be performed, with particular focus on evaluating accumulated neurotoxicity and changes in biological age during the two marginal decades.

The current formulation models Aβ dynamics within a given brain region. It does not explicitly model region-to-region spread of pathology (prion-like propagation) or network-level dysfunction (circuit disruption due to regional damage). These phenomena could be incorporated in future work by coupling multiple regional models through transport terms, but are beyond the scope of the present analytical framework, which focuses on establishing the local kinetic behavior and biological age concept within individual regions.

## Abbreviations

AD: Alzheimer’s disease
Aβ: amyloid beta
CV: control volume
F-W: Finke−Watzky
IDP: intrinsically disordered protein

## Acknowledgements

The support for this research was provided by the National Science Foundation (grant DMS-2451660) and the Alexander von Humboldt Foundation through the Humboldt Research Award.

## Supplemental Materials

## S1. Modeling senile plaque growth

This study follows the derivation developed in Kuznetsov (2025a), Kuznetsov (2025c). To simulate the growth of an Aβ plaque (illustrated in Fig. 1), the approach focuses on determining the total number of Aβ monomers, denoted as *N*, that accumulate within the plaque over time *t*. Doing so requires considering the average concentration of Aβ monomers deposited into the plaque, as proposed in Watzky *et al*. (2008):

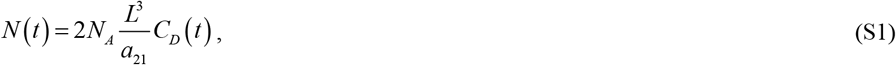

where *N*_*A*_ is Avogadro’s number. The factor of 2 on the right-hand side of Eq. (S1) accounts for the assumption that an Aβ plaque forms in the space between two adjacent neurons. Each neuron’s membrane releases Aβ monomers at a rate of *q*_*A*,0_, contributing symmetrically to plaque formation (Fig. 1).

An alternative approach, also based on Watzky *et al*. (2008), involves determining *N* (*t*) using the volume of an individual inclusion body, denoted as *V*_*ABP*_ :

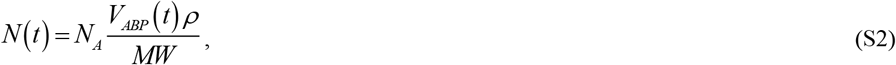

where *MW* represents the average molecular weight of an Aβ monomer.

Equating the right-hand sides of Eqs. (S1) and (S2) and solving for the volume of the Aβ plaque yield the following expression:

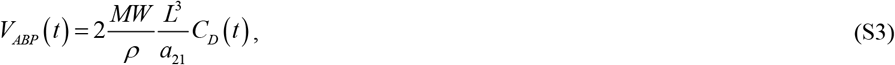

where 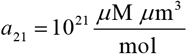 represents the conversion factor from 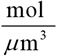 to *μ*M.

Assuming a spherical shape for the Aβ plaque, its volume is

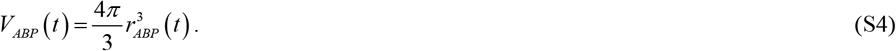

Equating the right-hand sides of Eqs. (S3) and (S4) and solving for the radius of the Aβ plaque yields the following expression:

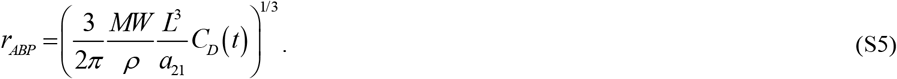

The size of an Aβ plaque is often quantified by its surface area, which can be measured from stained brain tissue sections examined under a microscope. Within the framework of this model, the plaque area can be calculated as follows:

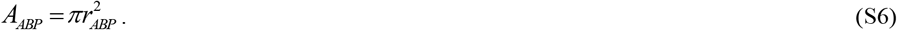

Aβ plaque load, defined as the percentage of the area occupied by amyloid plaques (Liu *et al*., 2017; Ikonomovic *et al*., 2008), can be determined using the following calculation:

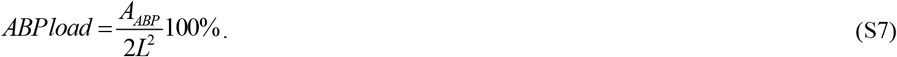

The factor of 2 in the denominator of Eq. (S7) accounts for the assumption in the model that a single plaque occupies two control volumes (Fig. 1).

## S2. Derivation of the “full analytical” solution for the scenario with constant, finite half-lives *T*1/ 2, *A, T*_1/ 2,*B*_, and *T*_1/2_,*D*

Summing Eqs. (23) and (24) gives:

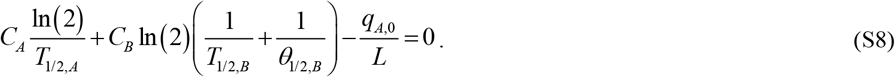

Solving for *C*_*A*_ yields:

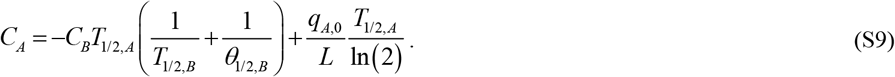

Substituting Eq. (S9) into Eq. (S8) results in a quadratic equation for *C*_*B*_ . The physically meaningful root, as concentration cannot be negative, is

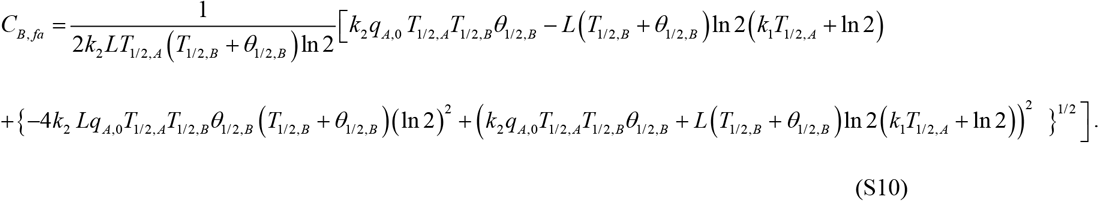

The subscript “*fa*” stands for “full analytical,” indicating that the analytical solution applies to finite, constant values of *T*_1/ 2, *A*_, *T*_1/ 2, *B*_, and *T*_1/ 2, *D*_ .

Substituting Eq. (S10) into Eq. (S9) and solving for *C*_*A*_ yield the following expression:

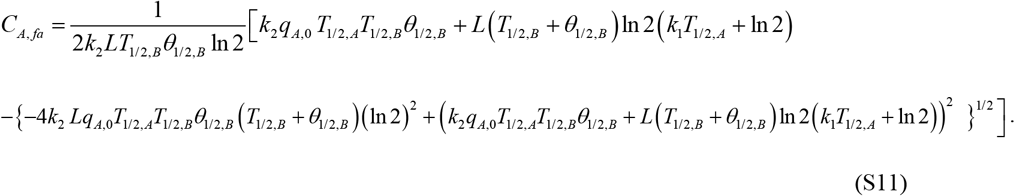

It is important to note that *C*_*B, fa*_ and *C*_*A, fa*_, as given by Eqs. (S10) and (S11), are independent of time.

Figs. S3b and S4b illustrate that the asymptotic values of *C*_*B*_ depend on both *T*_1/ 2, *A*,0_ and *T*_1/ 2, *B*,0_ . A similar dependence for *C*_*A*_ is observed in Figs. S3a and S4a. Notably, the numerical and analytical solutions exhibit excellent agreement once the time exceeds five years (Figs. S3 and S4).

Solving Eq. (6) for *C*_*D*_ for the case of constant, finite *T*_1/ 2, *D*_ value yields:

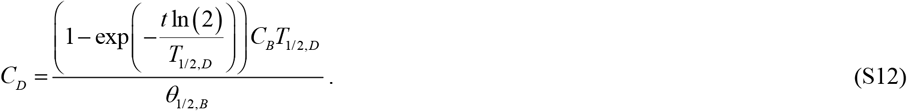

The value of *C*_*D, fa*_ (which is time-dependent) can be determined by substituting *C*_*B, fa*_ from Eq. (S10) into Eq. (S12). The plaque radius can be obtained by substituting *C*_*D, fa*_ from Eq. (S12) into Eq. (S5). Note that for small values of *t*, Eq. (S12) collapses to Eq. (21) when the expression from Eq. (17) is substituted for *C*_*B*_ .

Additionally, Eqs. (S12) and (17) can be used to predict the asymptotic value that *C*_*D*_ approaches as *t* →∞ :

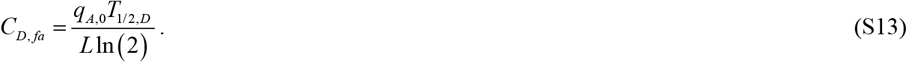

However, this asymptotic value is excessively large and unlikely to have biological significance. Utilizing Eq. (25) yields the following result:

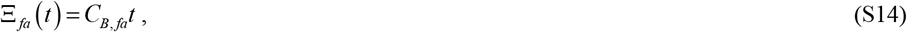

where *C*_*B, fa*_ is defined by Eq. (S10).

The biological age is calculated by substituting Eq. (S14) into Eq. (27):

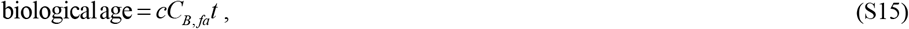

where the expression for *C*_*B, fa*_ is again provided in Eq. (S10).

By utilizing the “full analytical” solution from Eqs. (S15) and (S10), the following expressions are obtained for the dimensionless sensitivities of biological age to the half-lives of Aβ monomers and free aggregates, which reflect the efficiency of the Aβ degradation machinery:

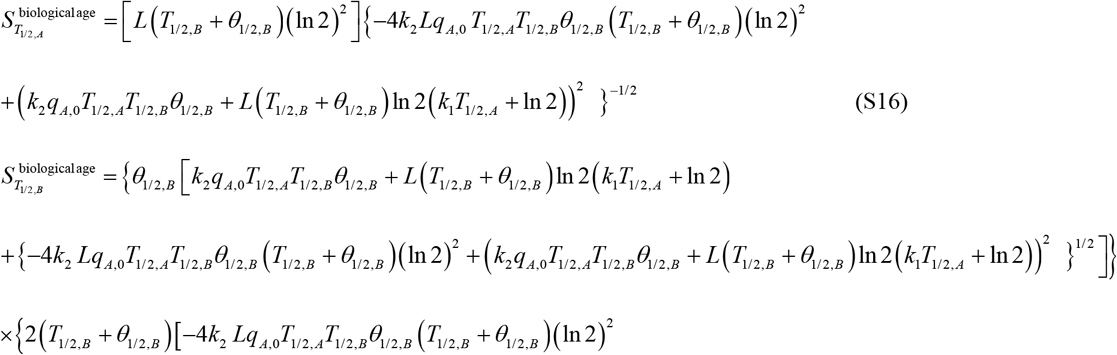

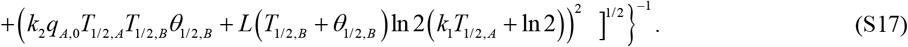

## S3. Numerical Solution

The system of differential equations in Eqs. (4)–(6), along with the initial conditions defined in Eq. (7), was solved numerically using MATLAB’s ODE45 solver (MATLAB R2024a, MathWorks, Natick, MA, USA). ODE45, which uses an adaptive timestep-step Runge-Kutta method, was chosen for its accuracy and robustness. To ensure high numerical precision, both the relative tolerance (RelTol) and absolute tolerance (AbsTol) were set to 1e-10.

## S4. Supplementary figures

**Fig. S1.**
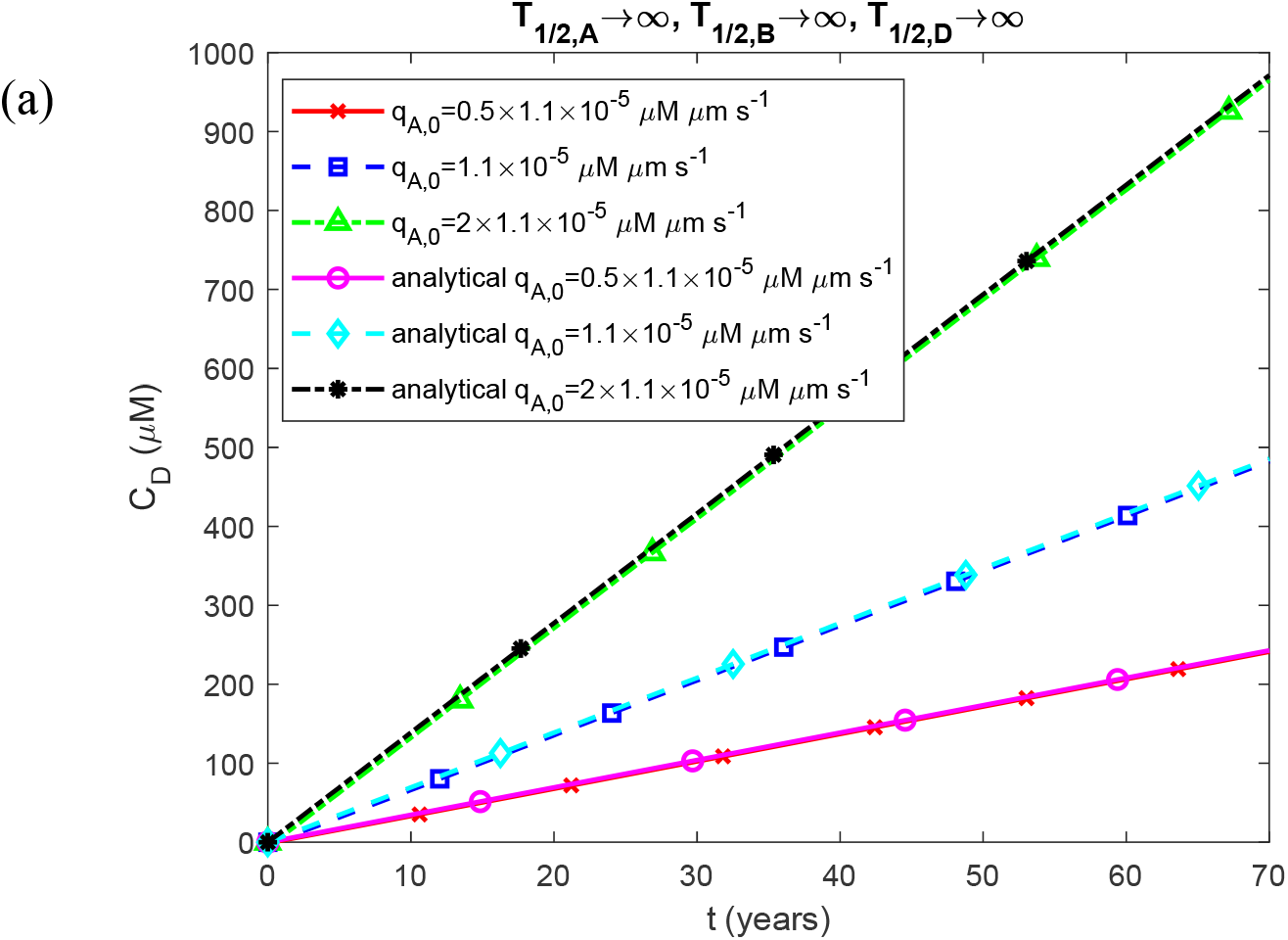
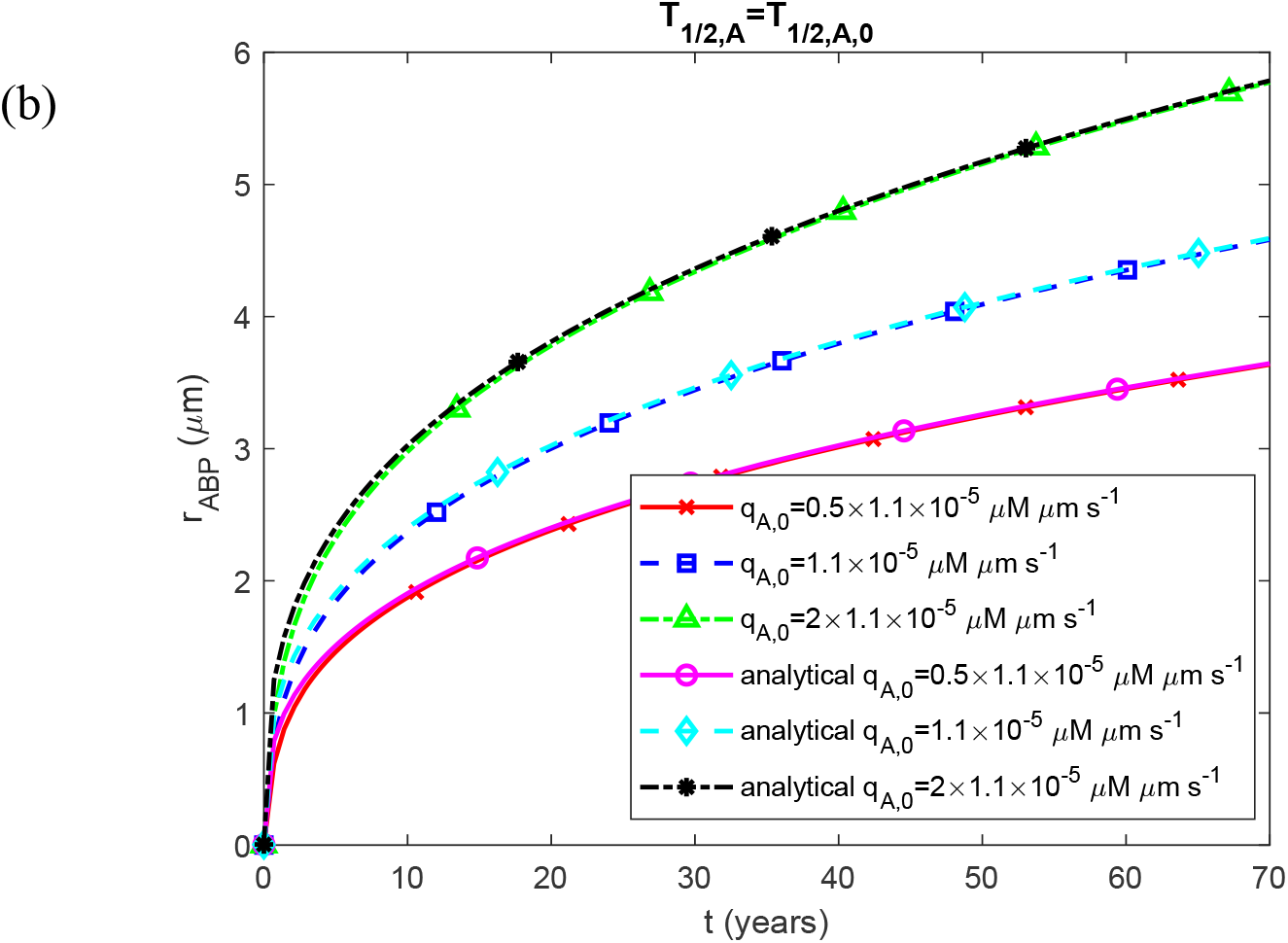
(a) Molar concentration of Aβ aggregates deposited in plaques, *C*_*D*_, as a function of time and (b) radius of a growing Aβ plaque, *r*_*ABP*_, as a function of time. The plots are presented for different values of Aβ monomer flux into the CV, *q*_*A*,0_ . Parameters: *T*_1/2, *A*_ =10^20^ s, *T*_1/2, *B*_ =10^20^ s, *T*_1/2, *D*_ =10^20^ s. The plotted analytical solutions are obtained from Eqs. (21) and (22).

**Fig. S2.**
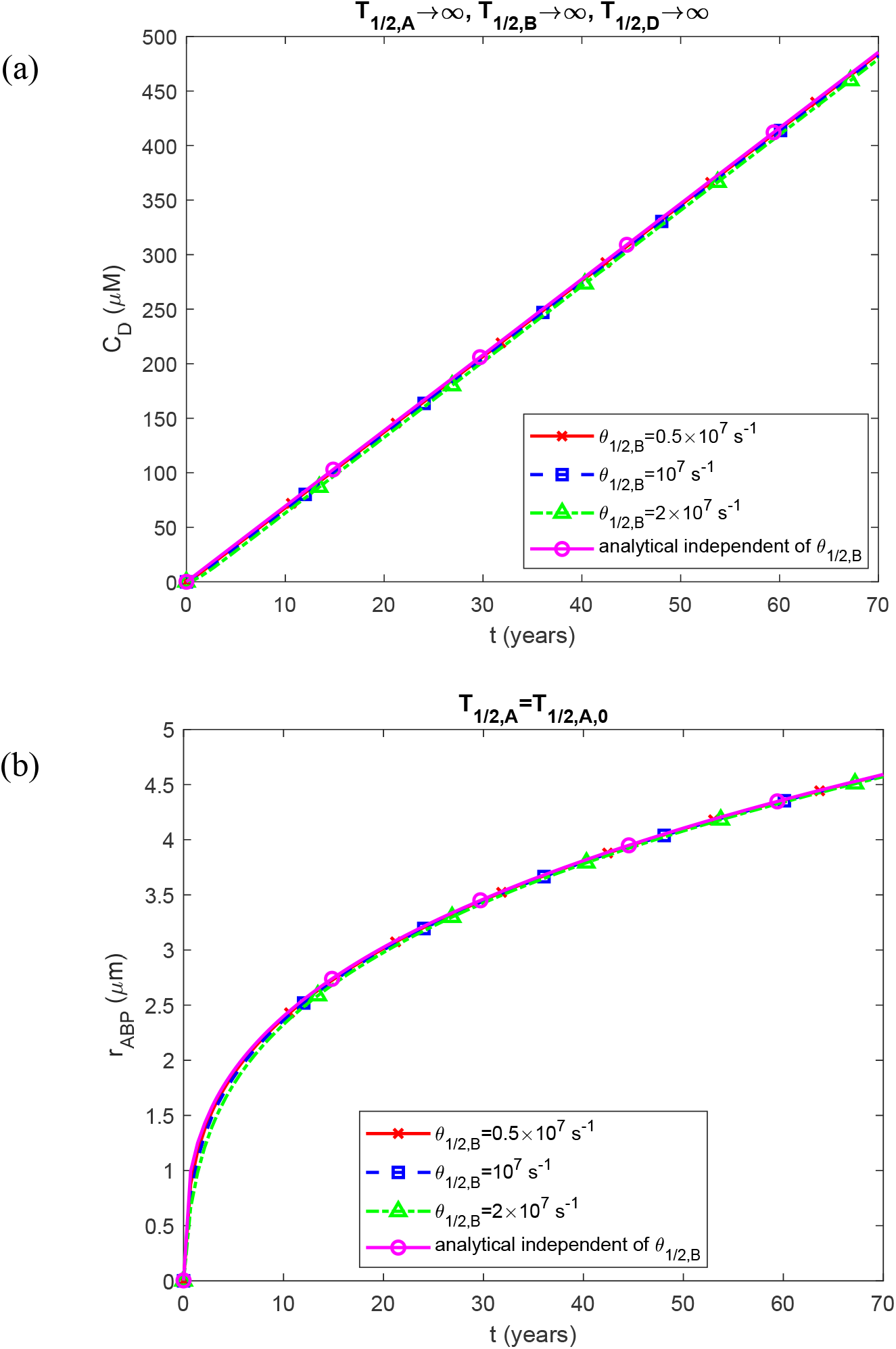
(a) Molar concentration of Aβ aggregates deposited in plaques, *C*_*D*_, as a function of time and (b) radius of a growing Aβ plaque, *r*_*ABP*_, as a function of time. The plots are presented for different values of half-deposition times of free Aβ aggregates into senile plaques, *θ*_1/2, *B*_ . Parameters: *T*_1/2, *A*_ =10^20^ s, *T*_1/2, *B*_ =10^20^ s, *T*_1/2, *D*_ =10^20^ s. The plotted analytical solutions are obtained from Eqs. (21) and (22).

**Fig. S3.**
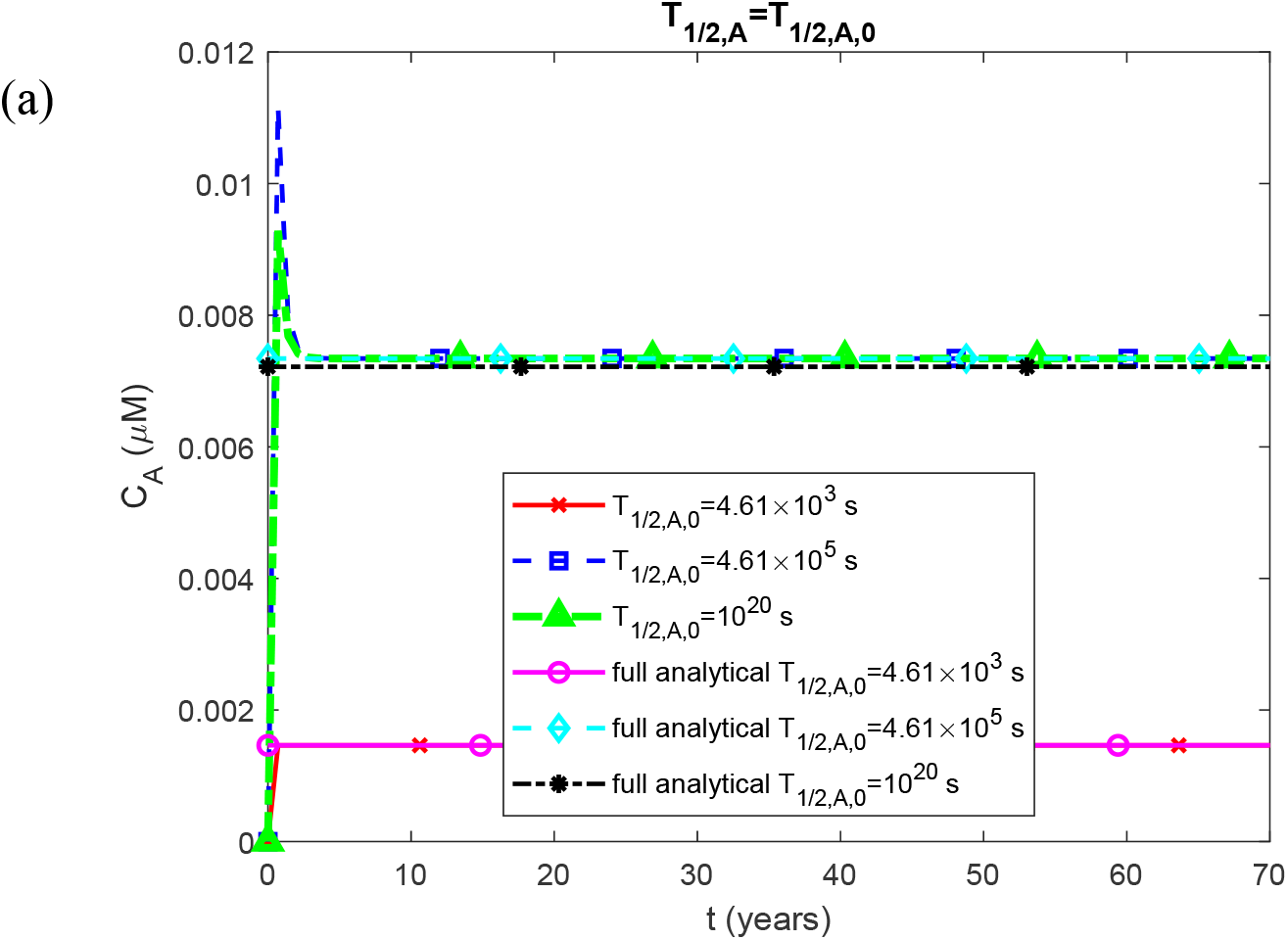
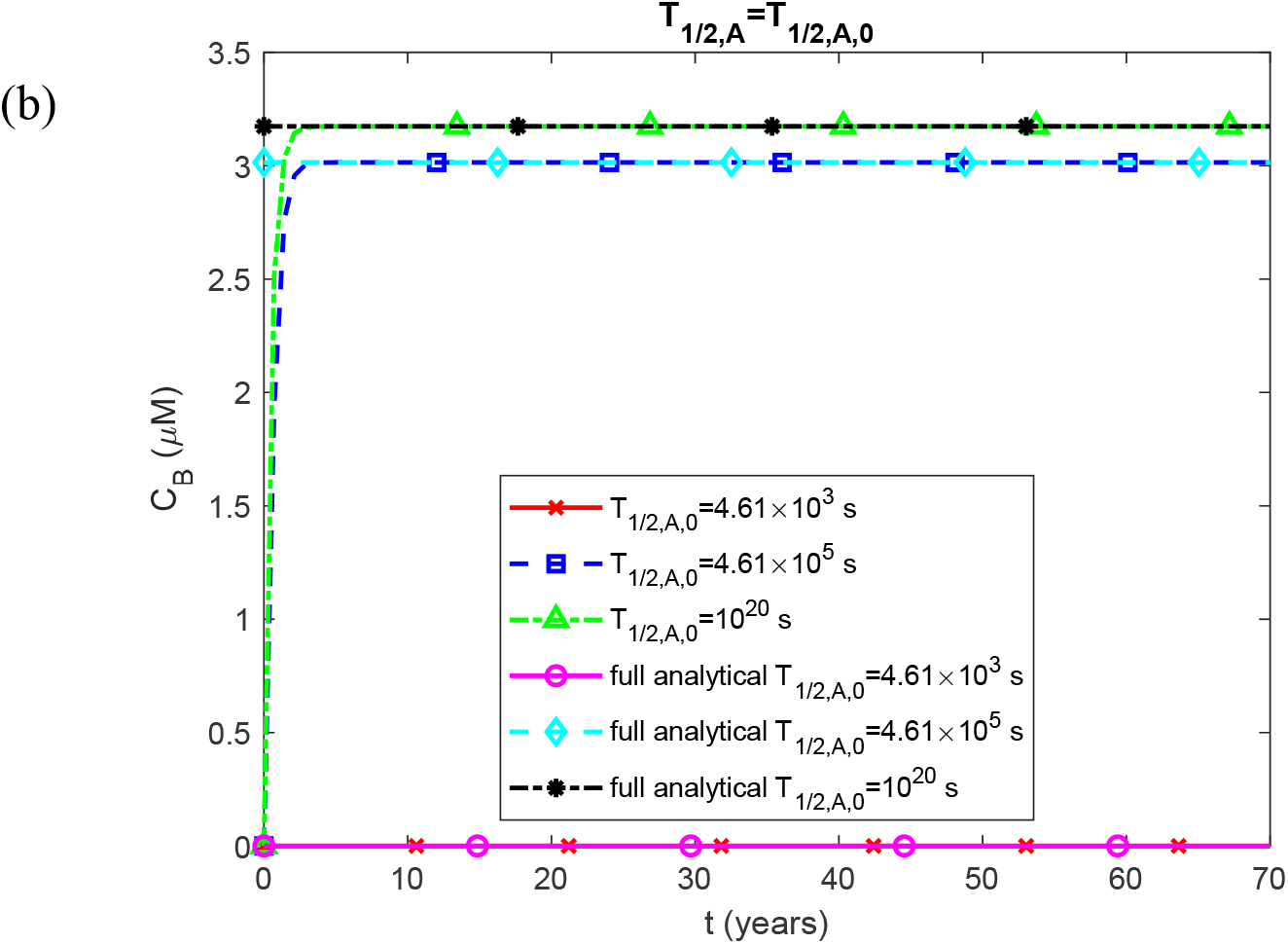
Molar concentration of (a) Aβ monomers, *C*_*A*_, and (b) Aβ oligomers, *C*_*B*_, as a function of time for different values of the half-life of Aβ monomers, *T*_1/2, *A*_ . Parameters: *T*_1/2, *B*_ =10^20^ s. The plotted “full analytical” solutions are derived from Eqs. (S11) and (S10). Note that in each plotted case *C*_*A*_ and *C*_*B*_ reach constant asymptotic values as time increases. This is essential for deriving the full analytical solution.

**Fig. S4.**
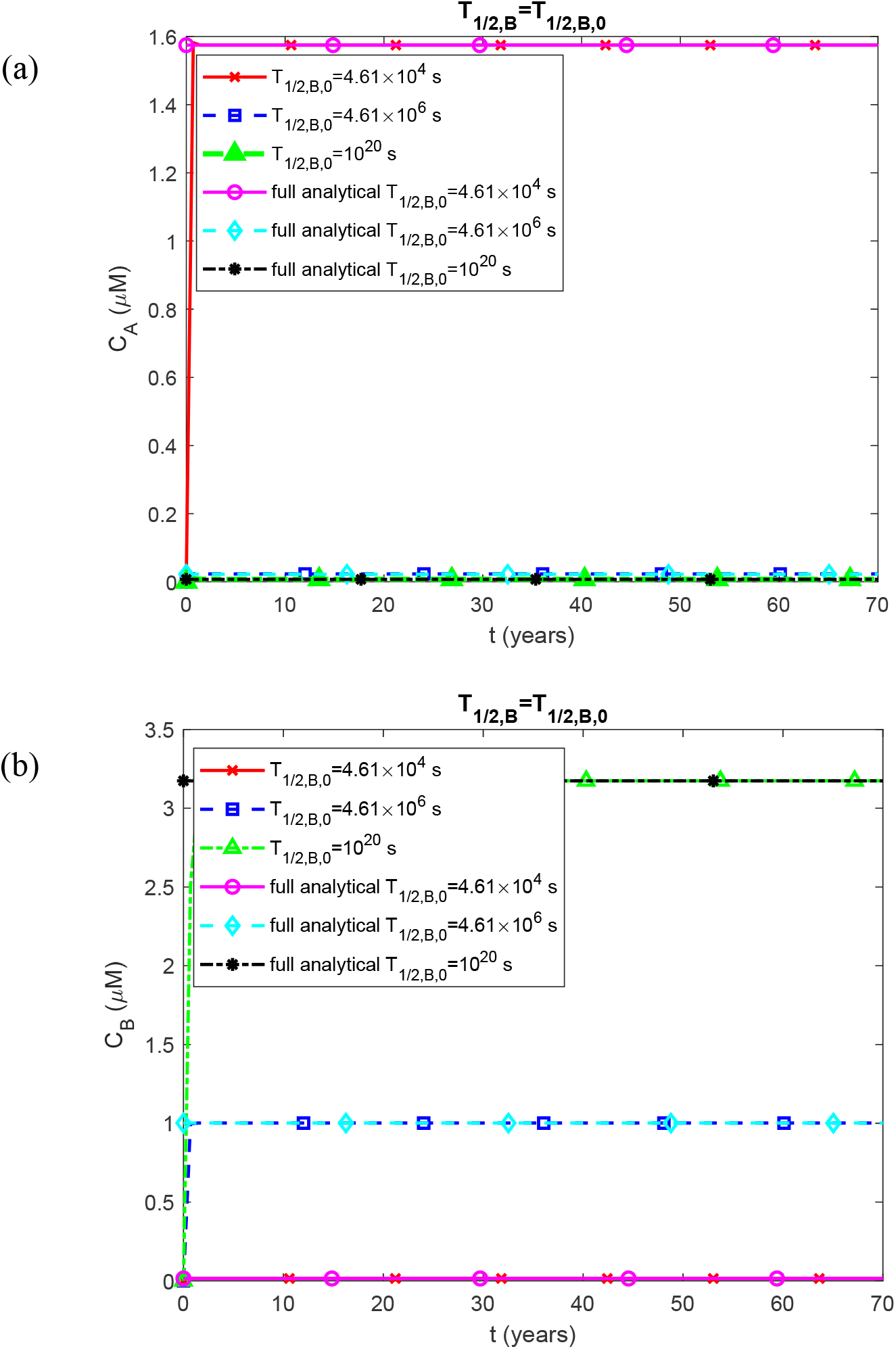
Molar concentration of (a) Aβ monomers, *C*_*A*_, and (b) Aβ oligomers, *C*_*B*_, as a function of time for different values of the half-life of Aβ oligomers, *T*_1/2, *B*_ . Parameters: *T*_1/2, *A*_ =10^20^ s. The plotted “full analytical” solutions are derived from Eqs. (S11) and (S10). Note that in each plotted case *C*_*A*_ and *C*_*B*_ reach constant asymptotic values as time increases. This is essential for deriving the full analytical solution.

**Fig. S5.**
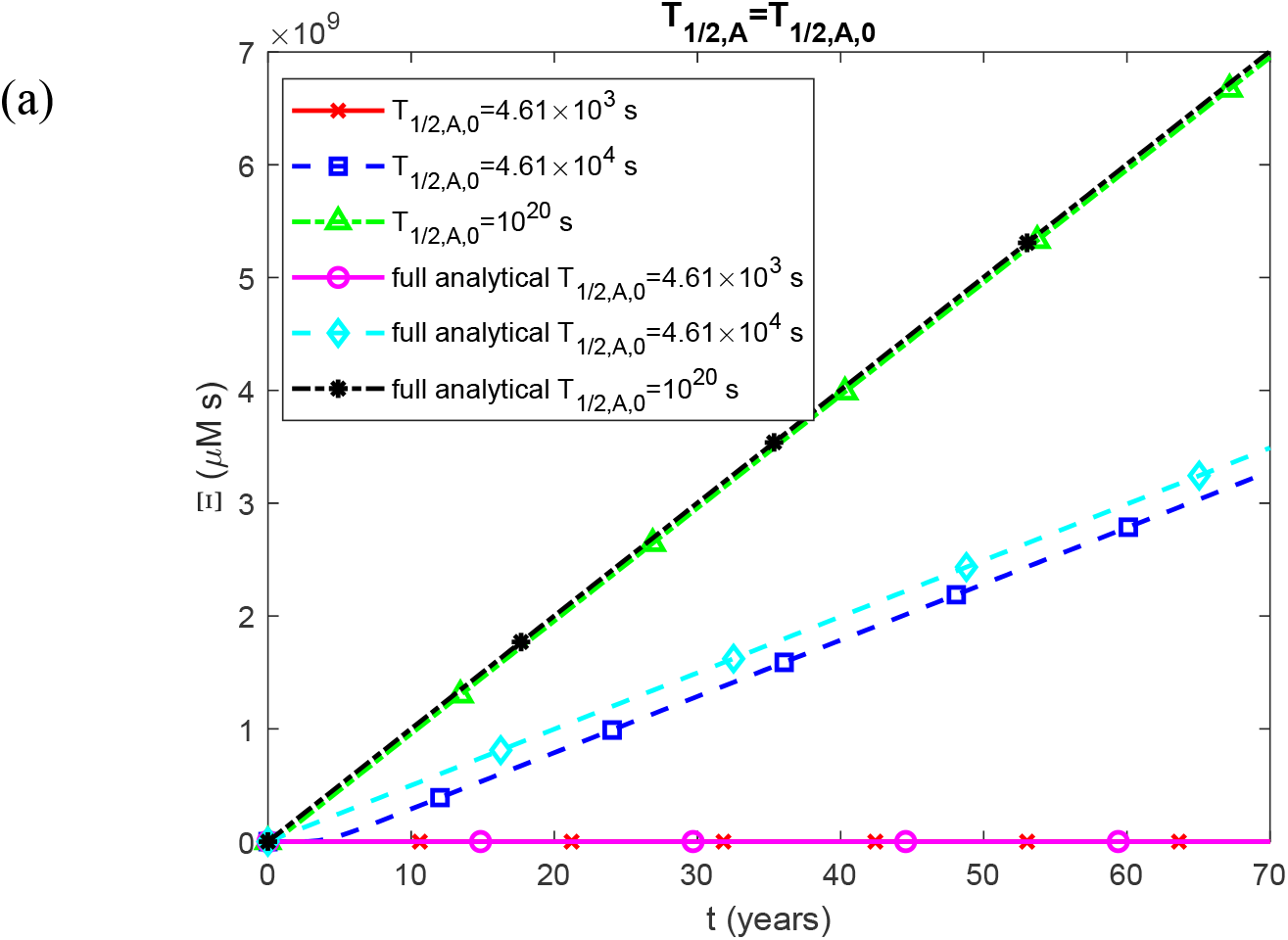
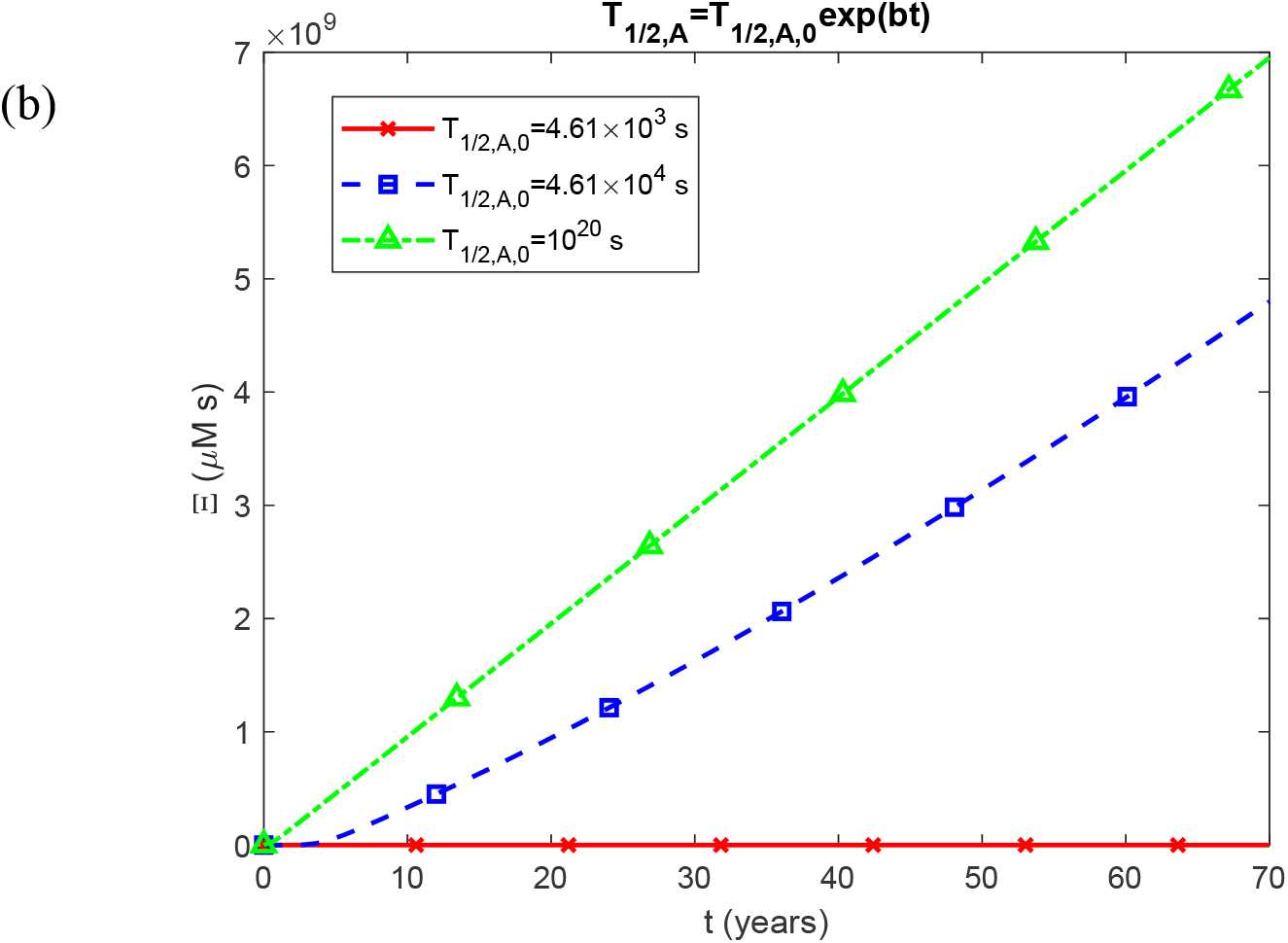
Accumulated neurotoxicity of Aβ oligomers as a function of calendar age. (a) The half-life of Aβ monomers, *T*_1/2,*A*_, remains constant regardless of calendar age. (b) The half-life of Aβ monomers, *T*_1/2,*A*_, increases exponentially with calendar age. Parameters: *T*_1/2, *B*_ =10^20^ s.

**Fig. S6.**
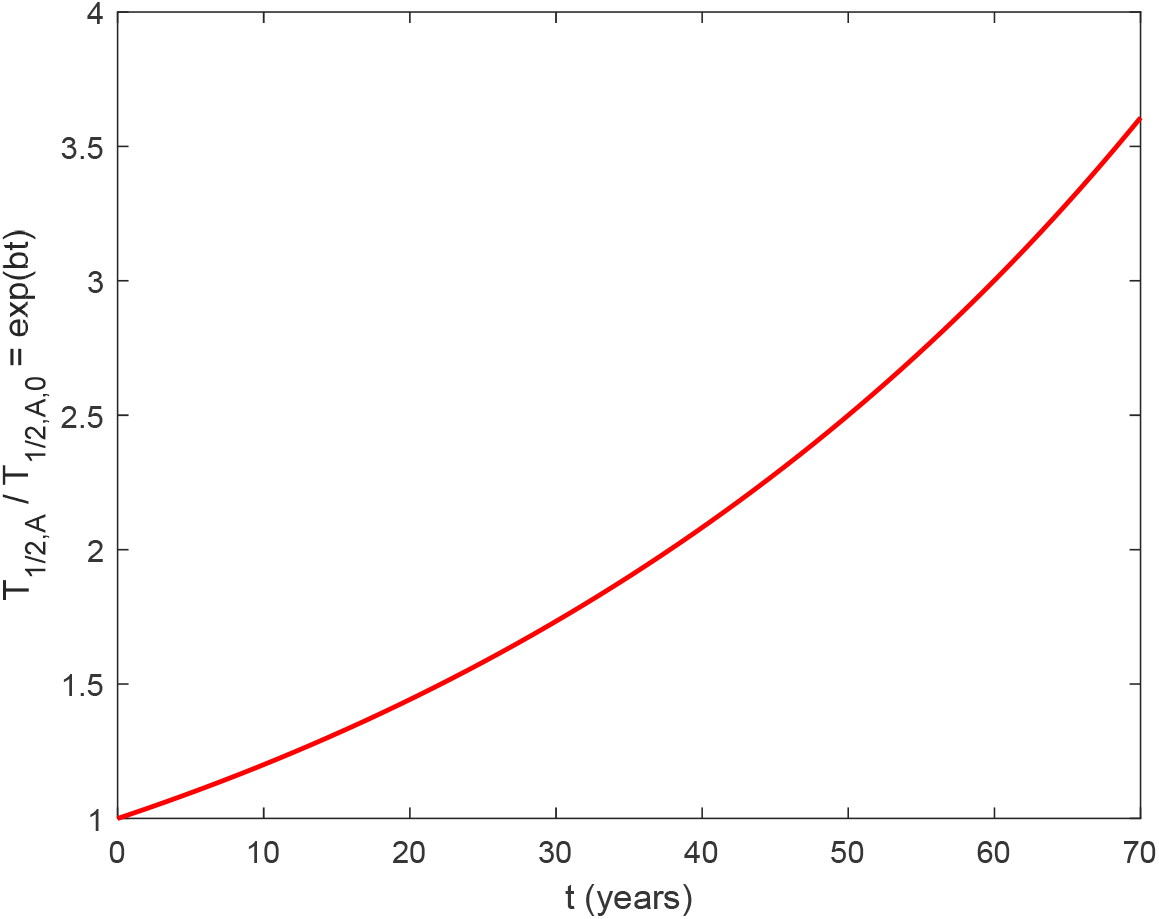
A function illustrating the exponential increase in the half-lives of Aβ monomers, *T*_1/2, *A*_ / *T*_1/ 2, *A*,0_ . In this scenario, the half-life of Aβ monomers is assumed to increase by a factor of 2.5 by the time the calendar age reaches 50 years, as described in Patterson *et al*. (2015). The same model is applied to the half-lives of free aggregates and aggregates deposited in Aβ plaques in the scenario of exponentially increasing half-life.

**Fig. S7.**
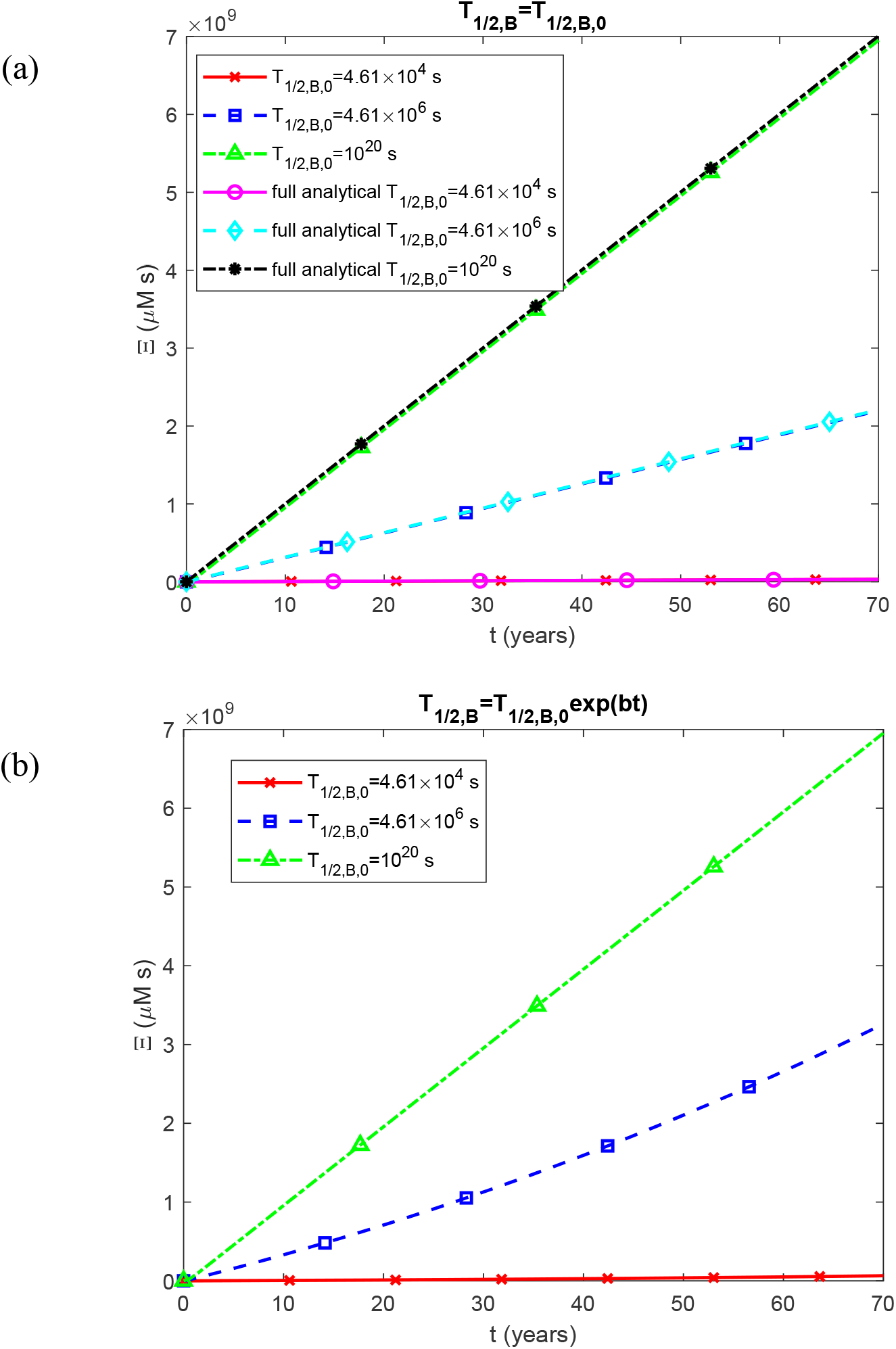
Accumulated neurotoxicity of Aβ oligomers as a function of calendar age. (a) The half-life of free Aβ aggregates, *T*_1/2,*B*_, remains constant over time. (b) The half-life of free Aβ aggregates, *T*_1/2,*B*_, increases exponentially with calendar age. Parameters: *T*_1/2, *A*_ =10^20^ s.

**Fig. S8.**
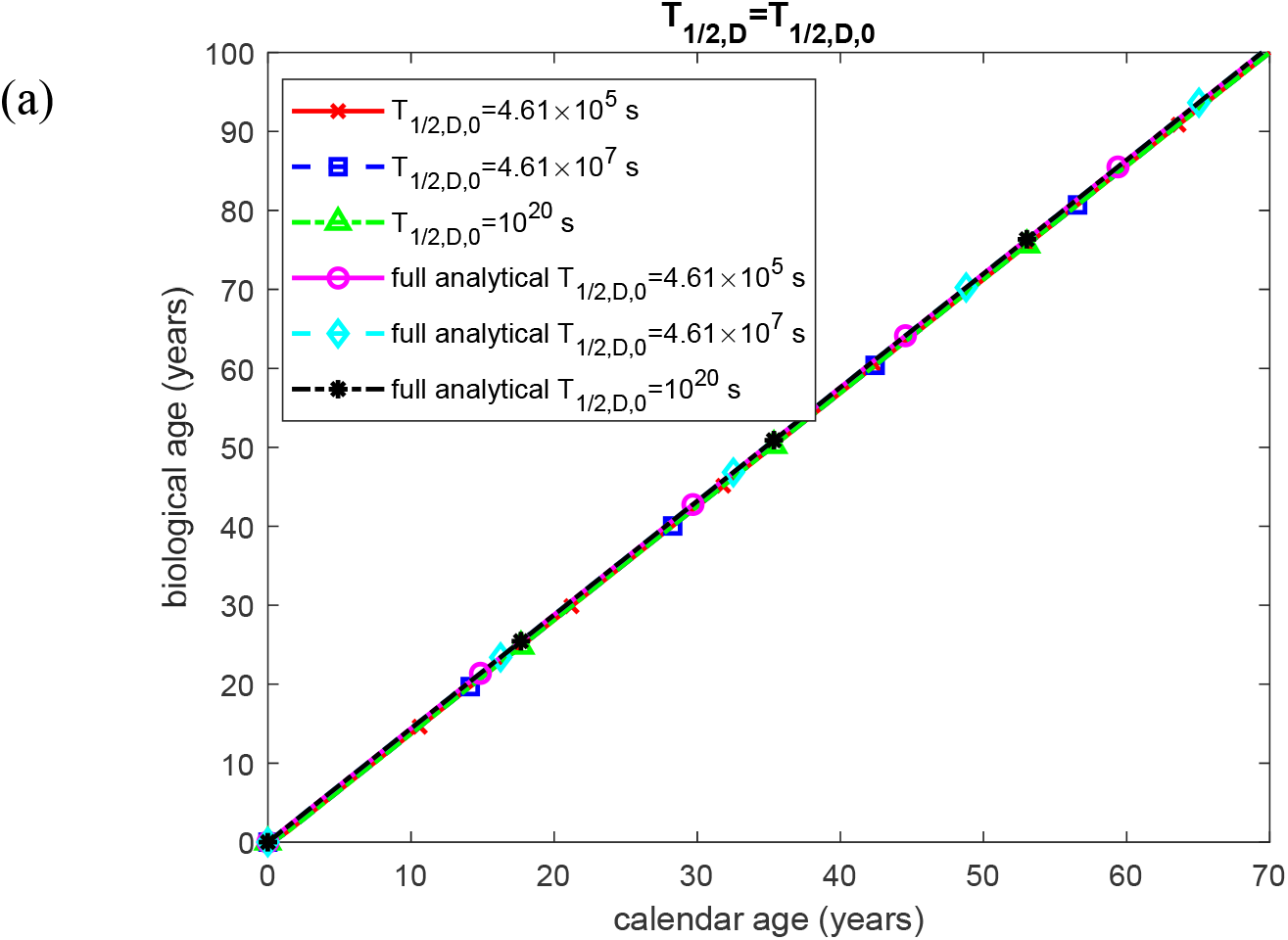
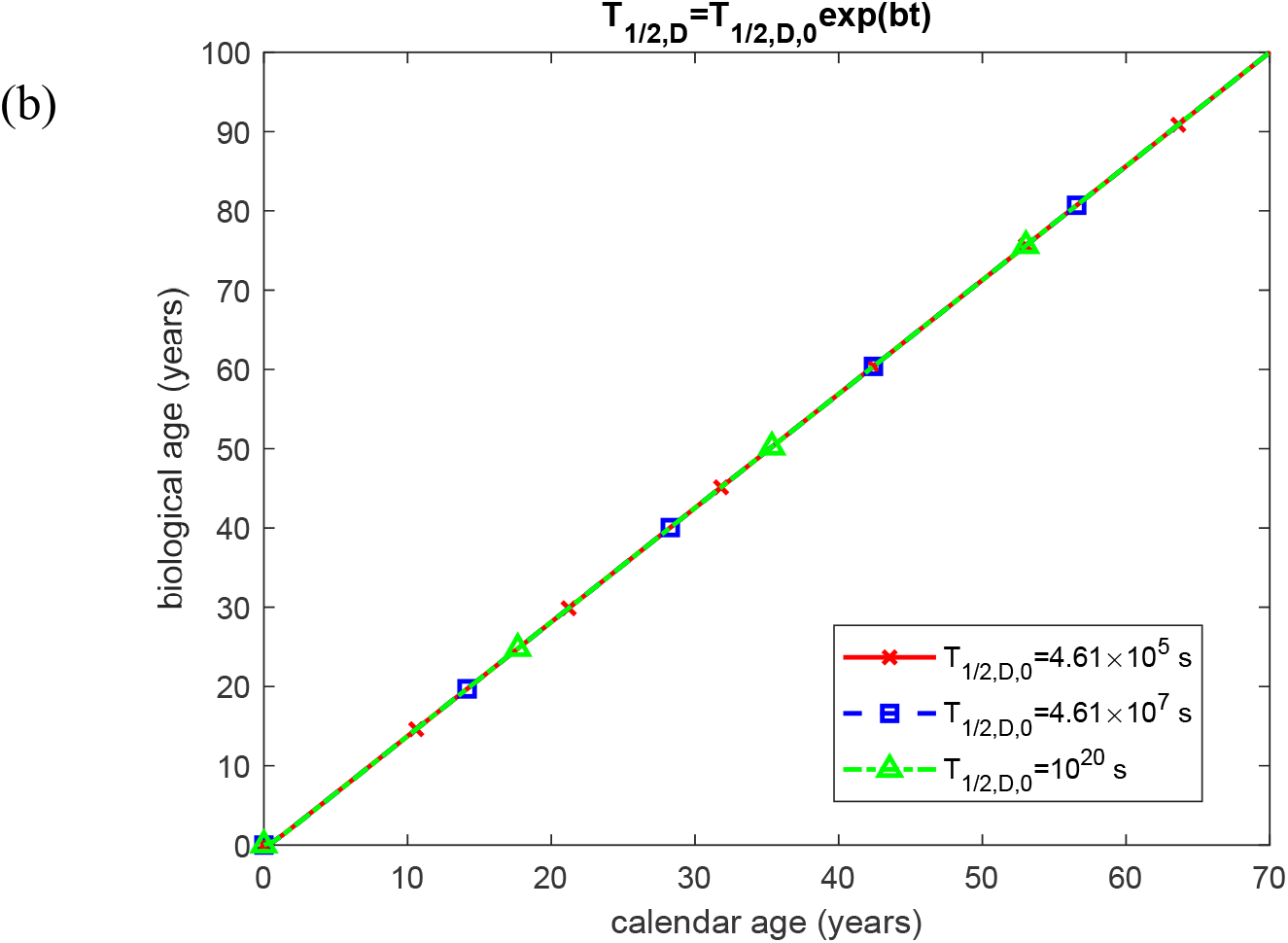
Biological age as a function of calendar age. (a) The half-life of Aβ aggregates after deposition into senile plaques, *T*_1/2,*D*_, remains constant over time. (b) The half-life of Aβ aggregates after deposition into senile plaques, *T*_1/2, *D*_, increases exponentially with calendar age. Parameters: *T*_1/2, *A*_ =10^20^ s, *T*_1/2, *B*_ =10^20^ s.

**Fig. S9.**
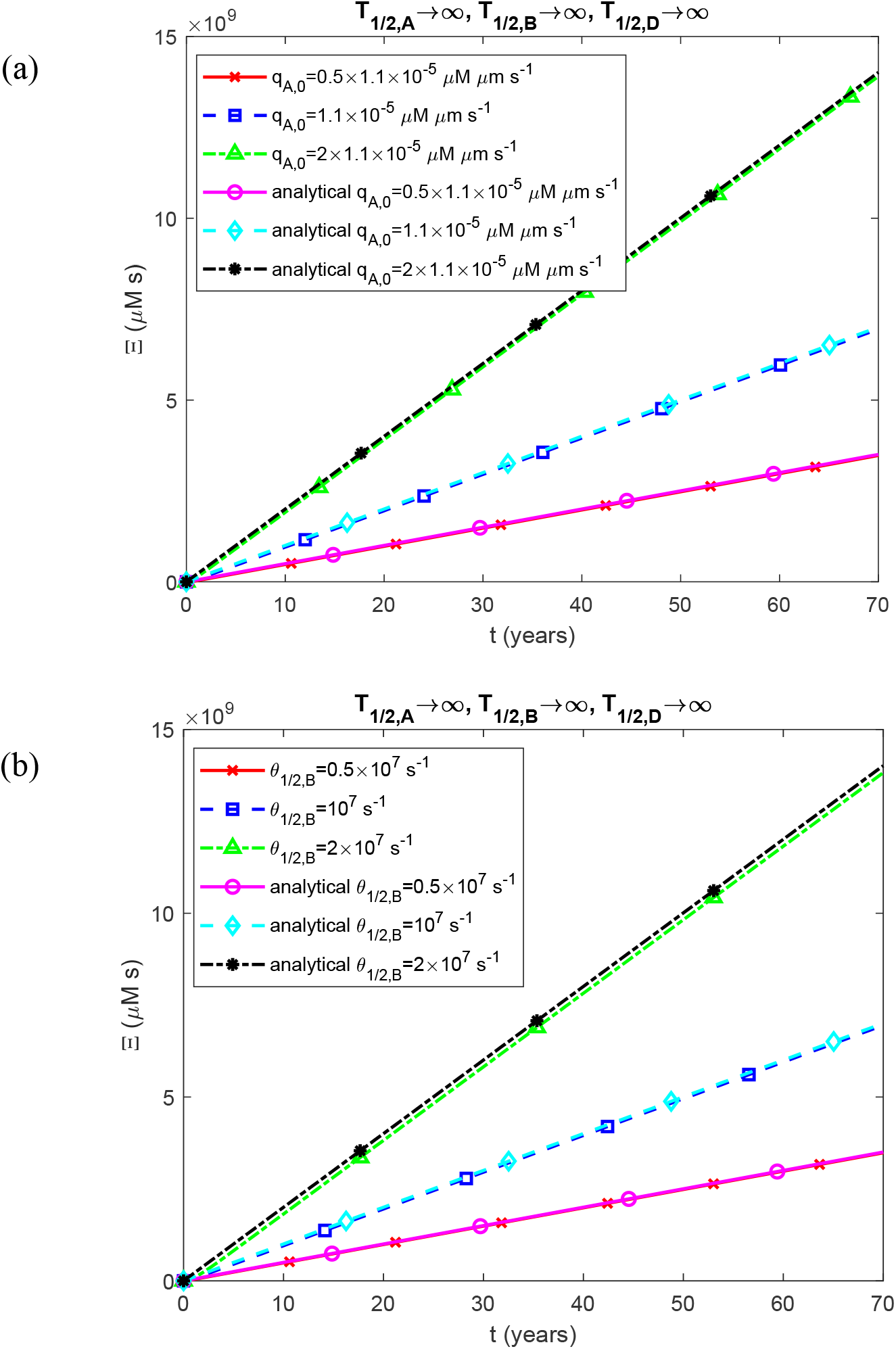
(a) Accumulated neurotoxicity of Aβ oligomers as a function of calendar age for different values of the flux, *q*_*A*,0_, of Aβ monomers per unit area at the left-hand face of the shaded CV shown in Fig. 1. (b) Accumulated neurotoxicity of Aβ oligomers as a function of calendar age for different values of the half-deposition time, *θ*_1/2, *B*_, of free Aβ aggregates into senile plaques. Parameters: *T*_1/2, *A*_ =10^20^ s, *T*_1/2, *B*_ =10^20^ s.

**Fig. S10.**
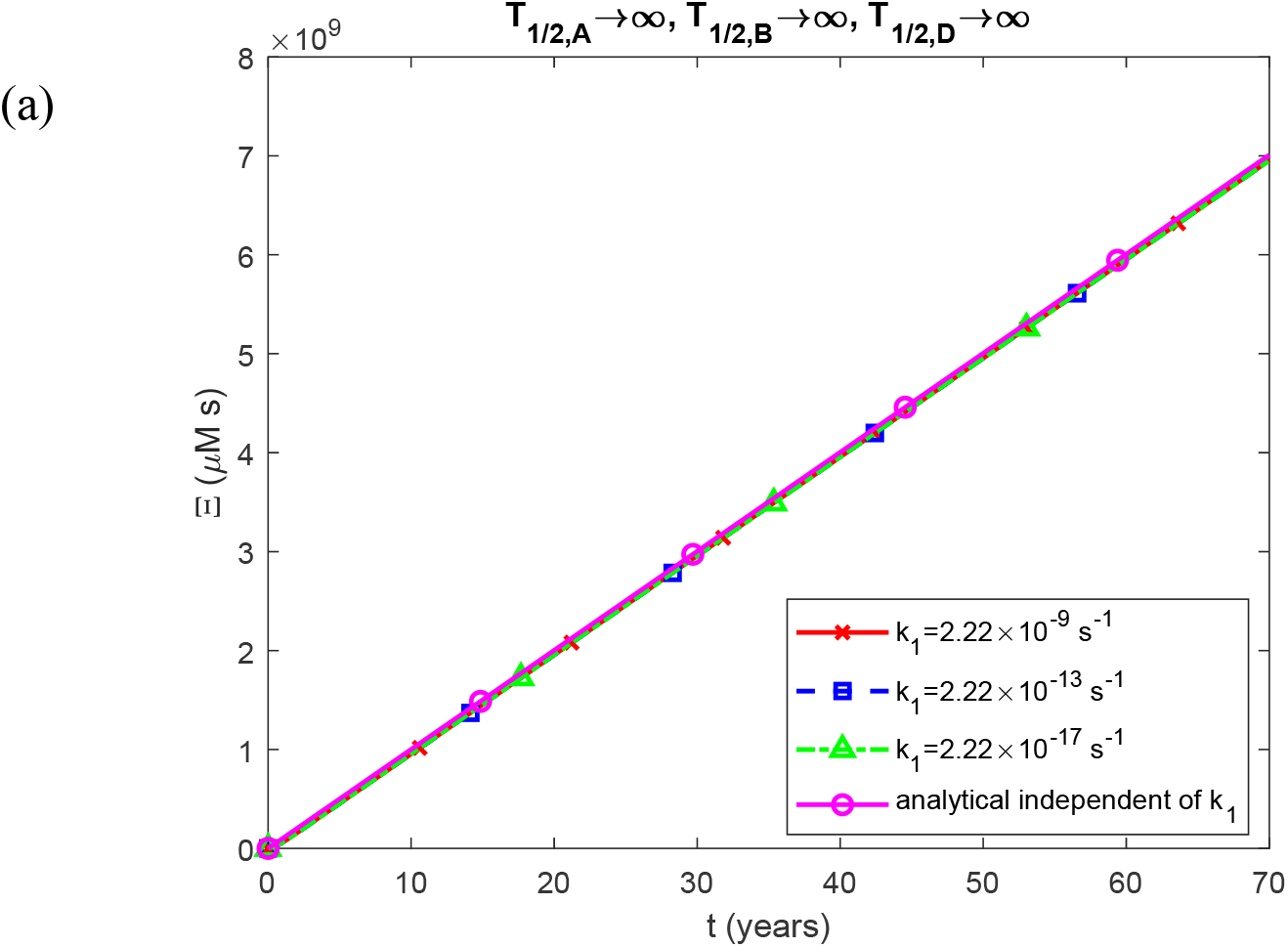
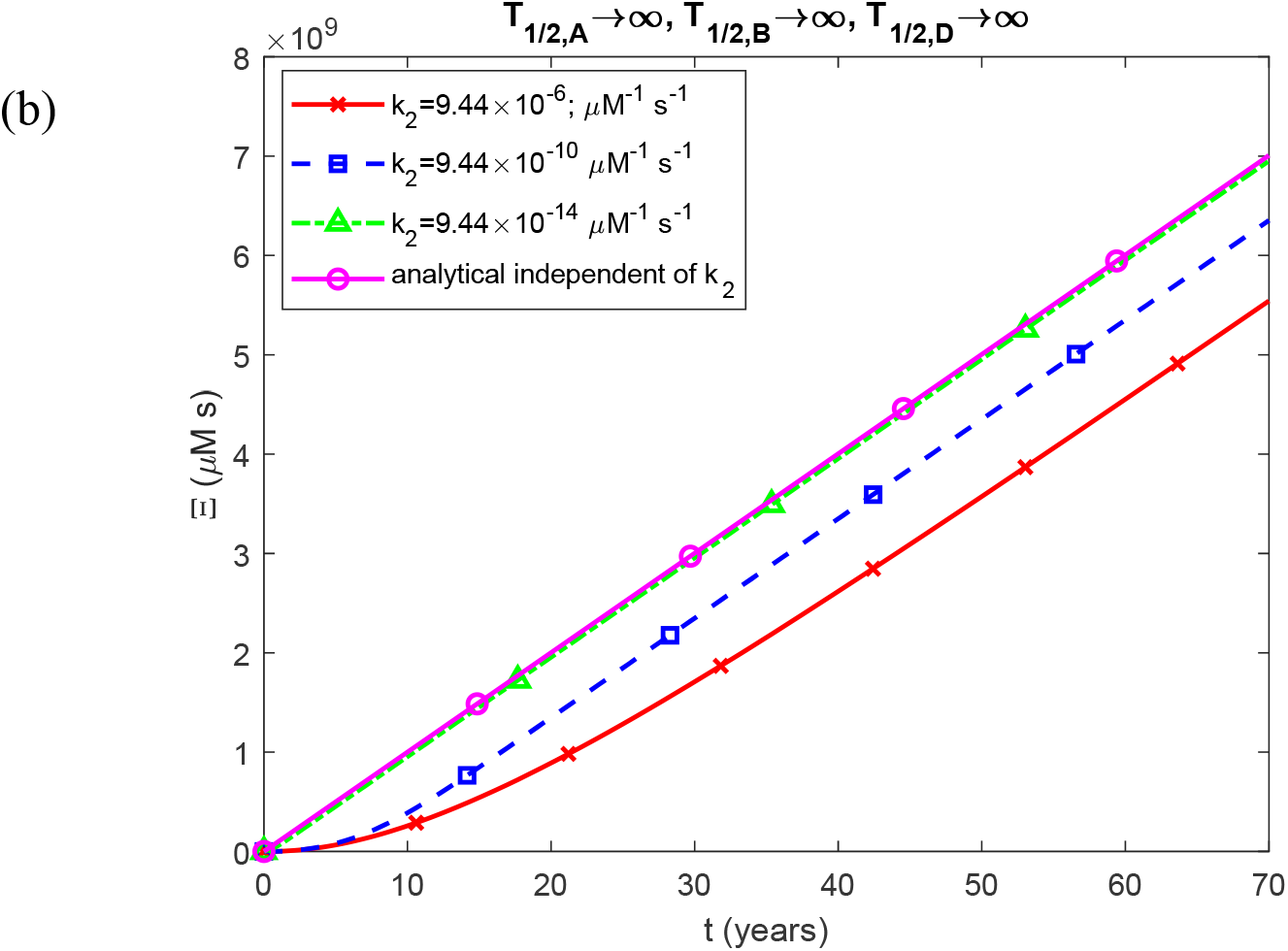
(a) Accumulated neurotoxicity of Aβ oligomers as a function of calendar age for different values of the nucleation rate constant, *k*_1_, in the first pseudo-elementary reaction step of the F-W model for Aβ peptide. Parameters: *k*_2_ = 9.44 × 10^−6^ μM^-1^ s^-1^, *T*_1/2, *A*_ =10^20^ s, *T*_1/2, *B*_ =10^20^ s, *T*_1/2, *D*_ =10^20^ s. (b) Accumulated neurotoxicity of Aβ oligomers as a function of calendar age for different values of the autocatalytic growth rate constant, *k*_2_, in the second pseudo-elementary reaction step of the F-W model for Aβ peptide. Parameters: *k*_1_ = 2.22×10^−9^ s^-1^, *T*_1/2, *A*_ =10^20^ s, *T*_1/2, *B*_ =10^20^ s, *T*_1/2, *D*_ =10^20^ s.

